# RNA binding proteins and glycoRNAs form domains on the cell surface for cell penetrating peptide entry

**DOI:** 10.1101/2023.09.04.556039

**Authors:** Jonathan Perr, Andreas Langen, Karim Almahayni, Gianluca Nestola, Peiyuan Chai, Charlotta G. Lebedenko, Regan Volk, Reese M. Caldwell, Malte Spiekermann, Helena Hemberger, Namita Bisaria, Konstantinos Tzelepis, Eliezer Calo, Leonhard Möckl, Balyn Zaro, Ryan A. Flynn

**Affiliations:** Stem Cell Program and Division of Hematology/Oncology, Boston Children’s Hospital, Boston, MA, USA; Department of Pharmaceutical Chemistry, Cardiovascular Research Institute, University of California, San Francisco, CA, USA; Max Planck Institute for the Science of Light, Staudtstr. 2, 91058 Erlangen, Germany; Department of Physics, Friedrich-Alexander-University Erlangen-Nuremberg, 91054 Erlangen, Germany; Department of Stem Cell and Regenerative Biology, Harvard University, Cambridge, MA, USA; Wellcome-MRC Cambridge Stem Cell Institute, University of Cambridge, Cambridge, UK; Department of Haematology, University of Cambridge, Cambridge, UK; Experimental Cancer Genetics, Wellcome Trust Sanger Institute, Hinxton, Cambridge, UK; Department of Biology, Massachusetts Institute of Technology, Cambridge, MA, USA; Koch Institute for Integrative Cancer Research, Massachusetts Institute of Technology, Cambridge, MA, USA; Harvard Stem Cell Institute, Harvard University, Cambridge, MA, USA

## Abstract

The composition and organization of the cell surface determine how cells interact with their environment. Traditionally, glycosylated transmembrane proteins were thought to be the major constituents of the external surface of the plasma membrane. Here, we provide evidence that a group of RNA binding proteins (RBPs) are present on the surface of living cells. These cell surface RBPs (csRBPs) precisely organize into well-defined nanoclusters that are enriched for multiple RBPs, glycoRNAs, and their clustering can be disrupted by extracellular RNase addition. These glycoRNA-csRBP clusters further serve as sites of cell surface interaction for the cell penetrating peptide TAT. Removal of RNA from the cell surface, or loss of RNA binding activity by TAT, causes defects in TAT cell internalization. Together, we provide evidence of an expanded view of the cell surface by positioning glycoRNA-csRBP clusters as a regulator of communication between cells and the extracellular environment.

## Introduction

Becoming a cell surface molecule is a highly regulated process, mainly accomplished by classical secretion through the ER and Golgi. These compartments are also where glycosylation occurs, which often goes hand in hand with secretion (Varki et al., 2022). While lipids and proteins have long been studied as glycoconjugates, we recently described RNA as a novel template for modification with complex glycans of the secretory pathways (Flynn et al., 2021). These “glycoRNAs” are presented on the external surface of living cells and have the ability to interact with immunomodulatory receptors like Siglec proteins (Flynn et al., 2021). The positioning of glycoRNAs on the cell surface raises many questions, including how these hybrid biopolymers are connected to the plasma membrane.

Numerous studies have focused on understanding the proteinaceous composition of the cell surface. Biochemical, enzymatic, and focused protein-sequencing efforts helped to establish the identity of cell surface receptors. With greater access to higher resolution, higher sensitivity, and lower cost mass spectrometry-based proteomics (MS), various methods have enabled an “unbiased” view of the cell surface proteome (Li et al., 2020). While the vast majority of characterized surface proteins harbor the expected transmembrane domains (or other biophysical connections to the lipid bilayer like GPI-anchor proteins (Varki et al., 2022)), there are reports of those lacking traditional surface protein signatures, in particular RNA binding proteins (RBPs). Nucleolin (NCL), a highly expressed nucleolar RBP important for ribosomal biogenesis, has been reported to be found on the cell surface since at least 1990 (Semenkovich et al., 1990). This observation has been reproduced across numerous biological contexts (Christian et al., 2003; Hovanessian et al., 2010), with some correlation between the presentation of cell surface Nucleolin (csNCL) and various cancer states (Brignole et al., 2021; Hovanessian et al., 2010; Joo et al., 2018). Further, csNCL has been connected to some viral entry mechanisms such as human respiratory syncytial virus (Tayyari et al., 2011). More recently, there have been other reports of RBPs as putative cell surface proteins (Didiasova et al., 2019; Yoshimura et al., 2021) however the extent and generality of this localization is unclear.

Organization of cell surface molecules can control features such as the biophysics and regulatory potential of the plasma membrane. Defining the spatial distribution of cell surface molecules and the relationship between them offers important insights into their mechanisms of action. With the advent of super resolution microscopy techniques, aspects of how growth factor receptors (Werbin et al., 2017), cell surface adhesion complex formation (Fischer et al., 2021), membrane protein stoichiometry (Douglass and Vale, 2005; Fricke et al., 2015), and cell surface protein heterogeneity (Owen et al., 2010) have been uncovered with unprecedented detail. Changes in the presence or physical configuration of these cell surface proteins can lead to cell state changes and this can be accomplished by various means: cell autonomous processes such as secreted proteases (Mentlein, 2004; Salmi and Jalkanen, 2005), liquid-liquid phase separation (Case et al., 2019; Sánchez and Tampé, 2023), and external factors like pathogenic enzymatic activity like viral sialidases (McAuley et al., 2019). In addition to state changes, cell surface remodeling can also modulate the binding of various extracellular molecules for eventual entry into the target cell.

Cell penetrating peptides (CPPs) are a class of molecules that interact with the cell surface and as their name suggests, can enter cells (Langel, 2022). The first CPP identified was a minimal domain of the Trans-Activator of Transcription (full length TAT) protein from human immunodeficiency virus 1 (HIV1). Its main function is to directly bind the Trans-activation response element (TAR) RNA to facilitate transcription of the HIV1 genome (Das et al., 2011; Kao et al., 1987). While other types of CPPs have been discovered (Langel, 2022), the TAT CPP (herein referred to as TAT) is the prototypical cationic CPP. TAT can traffic payloads into cells en masse and more broadly, “poly-basic” proteins like the arginine-rich region of TAT have similar abilities to bind and enter cells (Koren et al., 2011). Various mechanistic explanations have been proposed (Guo et al., 2016). Given the highly cationic nature of these domains, sulfated cell surface glycans have been implicated as a mechanism of cell surface adhesion (Rusnati et al., 1999; Tyagi et al., 2001; Urbinati et al., 2009). However, it is not yet understood if TAT may also functionally interface with cell surface sialoglycoRNAs (Flynn et al., 2021), which also carry negative charges.

Here we investigate the idea that mammalian cells present RBPs on the cell surface (cell surface RBPs, csRBPs). We provide evidence that RBPs are common components of the cell surface across a wide array of cell types. We verified this hypothesis through multiple orthogonal molecular and imaging strategies. Through super resolution reconstructions we define a tessellated pattern of cell surface domains of prototypic size. The csRBPs cluster with each other and away from the MHC-I complex. These domains contain RNA ligands and are in proximity to glycoRNAs. In addition, clustering is in part dependent on intact cell surface RNA. Finally, we show that the TAT-derived cationic cell penetrating peptide is functionally dependent on intact cell surface RNAs for their cellular uptake.

## Results

### Many canonical RNA binding proteins are present on the external surface of living cells

A small number of RBPs have been individually characterized as being presented on the surface of living cells. With proteome-scale datasets defining RBPs and advancements in cell surface enrichment technologies, we set out to establish the generality of this observation. To generate a database of RBPs, we aggregated RBPs detected across 48 datasets specifically characterizing RBPs (RBPomes, **Table S1**). We defined a protein as a high confidence RBP as one which is present in at least 11 RBP databases (see **Methods**). This yielded a list of 1072 RBPs (**Table S1**). We next defined cell surface localized proteins by intersecting eight, seven, and six cell-surface proteomes identified by biotinylation of lysines by sulfo-NHS-SS-biotin, periodate-mediated oxidation of glycans, and other methods, respectively (See **Methods**, **Table S1**, and **Figure 1A**). This results in a set of 6,237 proteins that have been identified on the cell surface across various biological conditions and cell types. For both of these analyses, we limited our datasets to human cells and tissues. We intersected the set of RBPs with the set of cell surface proteins to define a set of cell surface RBPs (csRBPs). We denote intracellular RBPs with their standard name (e.g. “DDX21”) and the cell surface presented form with the “cs” prefix (e.g. “csDDX21”). Examining the overlap, we found that 211 of 1,072 RBPs (19.6%) were identified on the cell surface using all three chemical approaches for cell surface tagging (**Figure 1B, Figure S1A**). If we require that an RBP exist in at least 5 of 21 cell surface proteomes, this results in 187 RBPs (17.4% of all RBPs, **Figure 1B**). This set of csRBPs were all found using periodate labeling and enrichment which is often used to capture sialoglycans (De Bank et al., 2003; Gahmberg and Andersson, 1977), suggesting that they are likely glycoproteins. Because not all glycans have sialic acids, we expanded our analysis to RBPs that appear in at least 5 of 21 surfaceomes agnostic to the method of collection and found a set of 293 high confidence csRBPs (**Table S1**). Requiring that an RBP only be identified with two of the surface labeling strategies resulted in 750 of 1,072 RBPs (69.9%) being classified as a more general set of csRBPs (**Figure 1B, Figure S1B**). Conversely, even across the large number of cell surface proteomes investigated, 161 RBPs were never found (15.0%, **Figure 1B**). Taken together, these findings suggest that RBPs are a significant and diverse component of the cell surface.

**Figure 1.**
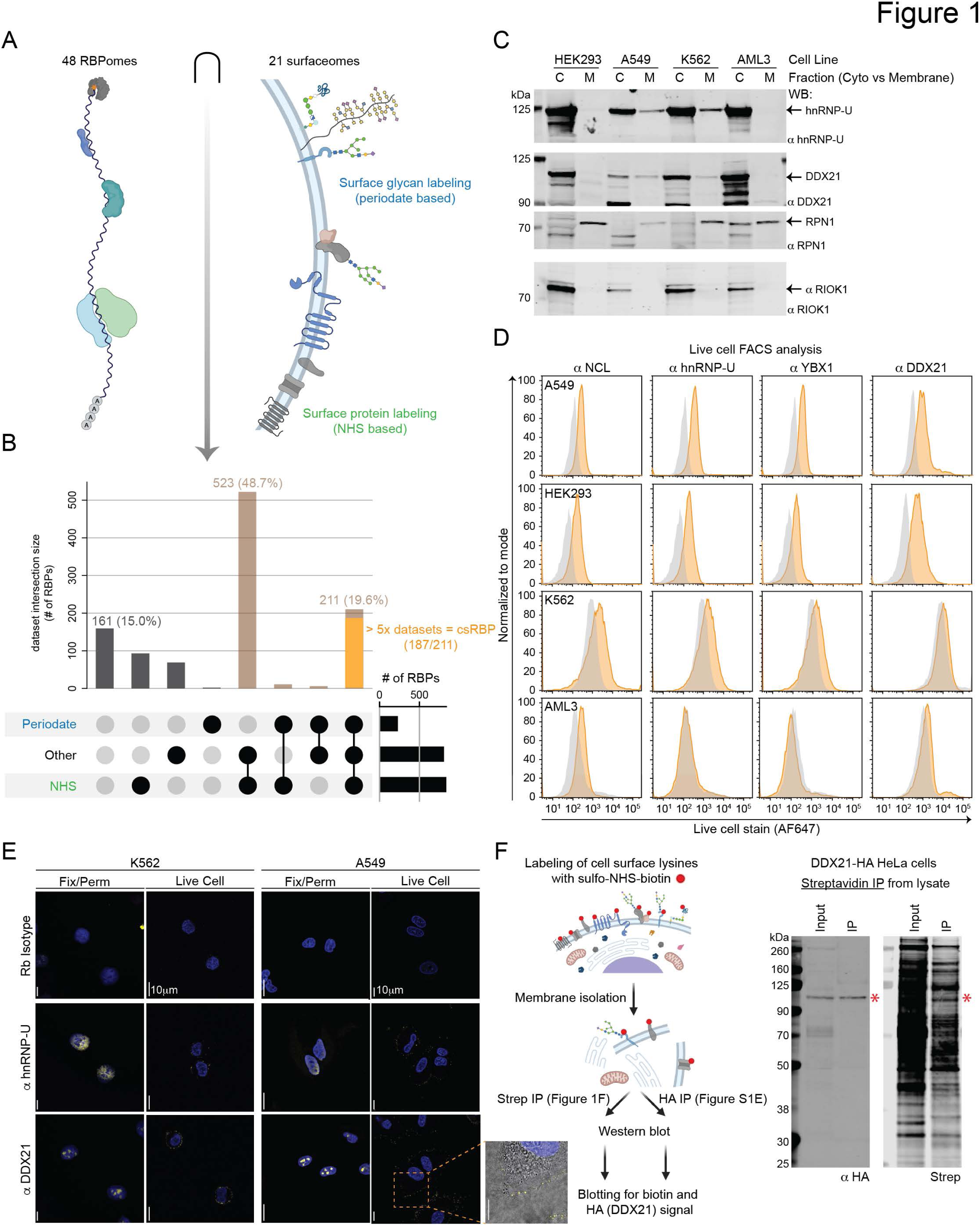
Expansive presentation of a select group of RNA binding proteins on the surface of living cells. A. Schematic of RNA binding proteins (RBPs, left) and methods to interrogate cell surface proteomes (surfaceome, right). B. Intersection analysis visualized using an upset plot examining RBPs identified as cell surface proteins stratified by different cell-surface-pulldown chemistries. The set size of each surfaceome is displayed as a bar plot at the right end of the intersection table. The right most intersection, RBPs found in all three classes of surfaceomes (cell surface RBPs, csRBPs), highlights the number of RBPs found in 5 or more datasets. C. Western blot analysis of cytosolic (cyto, C) and membrane (M) fractions from HEK293, A549, K562, and OCI-AML3 (AML3) cells. Equal cell fractions were loaded between C and M, and normalized across cell lines for the mass of protein in the C fraction. The expected molecular weight of the four proteins analyzed are noted with arrows and names of each protein and shown in kilodaltons (kDa). D. Flow cytometry analysis of the four cell lines in (C) stained live with antibodies targeting NCL, hnRNP-U, YBX1, DDX21 (yellow histograms), or isotype (gray histograms). E. Confocal microscopy of A549 and K562 cells. Cells were stain live (“Live Cell”) or after fixation and and permeabilization (“Fix/Perm”) with antibodies targeting DDX21, hnRNP-U, or isotype (yellow signal) and nuclei are stained with 4′,6-diamidino-2-phenylindole (DAPI, blue). Images were acquired using a 63x oil objective, a single z-slice is shown, and the scale bars are 10 µm. Zoomed region is also marked with a 10 µm scale bar and has the bright field signal overlaid to display the cell border. F. Cartoon of process to obtain cell surface labeled proteins for isolation and western blot detection (created with BioRender.com). Western blot analysis of HeLa cells expressing HA-tagged DDX21. Cells were labeled with sulfo-NHS-biotin before lysis, membrane organelles isolated, and biotinylated proteins enriched. Input lysate and enriched material was subjected to anti-HA (left) or streptavidin (right) blotting. The expected molecular weight for DDX21 is noted with a red asterisk.

To validate the putative csRBPs identified in the MS databases, we established a framework to these the hypothesis that each target is on the cell surface: a csRBP should fractionate with membrane organelles (e.g. plasma membrane), should be detectable using live cell flow cytometry and microscopy, and should be biochemically accessible to cell surface labeling reagents. We first performed a crude biochemical fractionation of cells, separating cytosol from bulk membrane organelles (a mix of ER, Golgi, mitochondria, and plasma membranes among others (Flynn et al., 2021)). We performed these experiments in four cell lines, two adherent (HEK293 and A549) and two suspension (K562 and OCI-AML3 (abbreviated AML3)). Western blot analysis of a known endoplasmic reticulum (ER) protein RPN1 and cytosolic RBP (RIOK1, an RBP never found in cell surface proteome datasets) demonstrated successful fractionation (**Figure 1C**). Two putative csRBPs DDX21 and hnRNP-U revealed strong signal in the cytosolic fraction and specific accumulation in the membrane fractions (**Figure 1C**), except from the OCI-AML3 cell line, which had no detectable hnRNP-U or DDX21 in the membrane fractions by Western blotting. Notably, despite RIOK1’s relatively high cytoplasmic abundance, the protein was largely undetectable in membrane fractions, supporting the predictive power of the csRBP list. To test if the presence in the membrane fraction could be attributed to cell surface localization, we stained living cells with commercially available, widely validated antibodies targeting four putative csRBPs (NCL, hnRNP-U, YBX1, and DDX21) and performed flow cytometry analysis (**Figure S1C**). All four cell lines exhibited surface presentation of at least one putative csRBP, and we found that levels of csRBP expression vary between cell lines (**Figure 1D**). Consistent with the lack of csRBPs appearing in the OCI-AML3 membrane fraction, OCI-AML3 cells stain weakly above isotype control, suggesting this cell line may express a different or less diverse suite of csRBPs.

We next assessed the spatial distribution of putative csRBPs. Cells were fixed and permeabilized (**Methods**) prior to staining with antibodies targeting DDX21 and hnRNP-U. By confocal microscopy we validated that antibodies targeting DDX21 and hnRNP-U generated nucleolar and nucleoplasmic signal, respectively. This distribution is consistent with the known localization of these proteins (**Figure 1E, Figure S1D**). We next performed live-cell staining, again using the anti-DDX21 and anti-hnRNP-U antibodies. Both DDX21 and hnRNP-U antibody staining manifested as clusters on the surface of cells (**Figure 1E**). These clusters were evident on A549 (adherent) and K562 (suspension) cells, although the clusters on A549 cells appear less densely distributed and more numerous (**Figure 1E**).

While these antibody-based and MS datasets (**Figure 1A**) support the cell surface presentation of RBPs like DDX21 and hnRNP-U, these experiments fail to assess whether we are detecting peptide fragments or full-length RBPs on the cell surface. To address this, we took a sequential, biochemical approach. We selectively biotinylated the cell surface of HeLa cells expressing DDX21 with a C-terminal HA-tag (Calo et al., 2015) using sulfo-NHS-biotin. We employed the HA-tag to enable higher specificity of detection by western blot. Crude membrane fractions were isolated as before and from these cells after surface labeling after which streptavidin capture was performed. Analysis of an anti-HA western blot generated from the streptavidin IP showed the selective enrichment of full length DDX21 protein (**Figure 1F**, left). Detection of biotinylated products also revealed a band at the expected molecular weight of full length DDX21 (**Figure 1F**, right). Reversing the capture order (anti-HA immunoprecipitation (IP) first) similarly demonstrated capture of full length DDX21. A weak but specific biotinylated band was enriched via the HA antibody at the expected molecular weight of DDX21 (**Figure S1E**). The overlap of biotin signal with anti-HA signal at DDX21’s expected molecular weight suggests that, at least in the case of DDX21, full-length csRBPs are able to be presented on the surface of living cells. Finally, to understand if csRBP deposition on the cell surface occurs in cis or in trans (perhaps from lysed cells or media contamination) we performed a co-culture experiment. Growing OMP25-eGFP (mitochondria) tagged HeLa cells with the DDX21-HA tagged HeLa cells and subsequently staining for cell surface HA signal, we found that GFP positive cells had little to no surface HA signal while GFP negative cells had clear cell surface HA puncta (**Figure S1F**), as we saw when staining with the endogenous anti-DDX21 antibody (**Figure 1E**). Taken together, these data support a model where cells expose RBPs to the extracellular environment.

### csRBPs are abundant and common members of cell surface proteomes

Aggregating across labs, species, cell types, processing methods, and Mass Spec methods (**Figure 1**) demonstrated the ubiquity of observing csRBPs; however, the heterogeneity of data makes quantitative comparisons more challenging. To rigorously examine the cell surface proteome and to directly validate publicly available data, we generated a new set of cell surface proteomics data. To ensure robust detection of only topologically extracellular proteins, we used a two-step selection process. First we labeled living cells with sulfo-NHS-SS-biotin (S-S; disulfide link) which should be cell impermeable. Second, we performed biochemical isolation of crude membranes (plasma membrane, ER, Golgi, mitochondria, etc) to separate highly abundant nuclear and cytosolic proteins (**Figure 2A, Figure S2A**). We performed this on four cell lines: HEK293, A549, OCI-AML3, and K562. We generated datasets of proteins detectable from bulk cytosol, bulk membrane, biotin-enriched cytosol, and biotin-enriched membrane (Mem-IP) fractions. After assessing and confirming that the biological triplicates from each sample type demonstrated strong correlation with each other (**Figure S2B**), we performed GO term analysis of proteins from the Mem-IP of the four cell lines. Enriched GO terms for cellular compartment (CC) indicated isolation of proteins associated with membranes or the cell surface (**Figure 2B**), confirming that our paired chemical and biochemical cell surface proteome labeling method targets cell-surface proteins with high specificity. When examining the biological process and molecular function GO terms, we found strong enrichment for terms related to RNA processes (**Figure 2B, S2C**).

**Figure 2.**
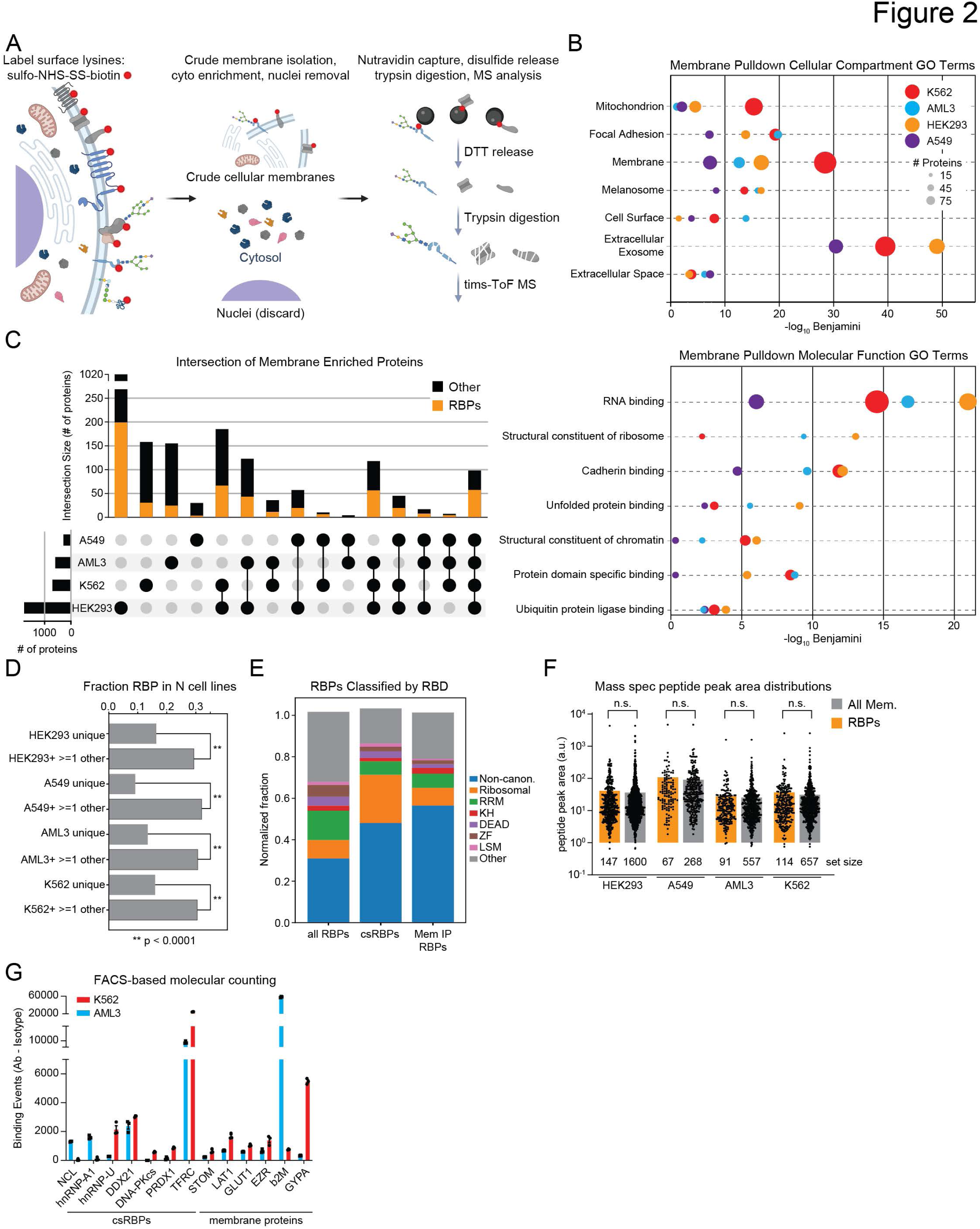
csRBP presentation is at parity with other canonical surface proteins. A. Experimental strategy to first label cell surface proteins while they are alive with a biotinylation reagent that can be cleaved by reduction. Subsequently, nuclei are isolated and discarded, and soluble cytosolic components are separated from membranous organelles. Input and biotin enriched proteins are then subjected to trypsin digestion and identification using a high resolution tims-ToF mass spectrometer (MS). Created with BioRender.com B. Gene ontology (GO) cellular compartment (CC) analysis of membrane enriched (Mem-IP) proteins identified by MS from HEK293, A549, K562, and OCI-AML3 (AML3) cells. The top 5 CC terms across the four cell lines were intersected and the union is displayed with the number of proteins in each term for each cell line (colors) represented by circle size and plotted on the x-axis by their significance. C. Intersection analysis visualized using an upset plot examining all Mem-IP hits across the four cell lines. The set size of each Mem-IP dataset is displayed as a bar plot at the left end of the intersection table. For each bar, the number of RBPs is overlaid in orange. D. Bar plot analysis of fraction of csRBPs found as uniquely present in one of the four cell lines (e.g. HEK293 unique) as compared to csRBPs found in at least two of the cell lines (e.g. HEK293+ >= 1 other). Statistical differences were evaluated with Chi-squared test. ** p < 0.0001. E. RNA binding domain (RBD) analysis of all RBPs, csRBPs defined in Figure 1, and csRBPs identified in the Mem-IP hits (Mem IP RBPs). Each class of RBD is represented with a different color in the stack bar plot. The sum of each class is > 1 because some RBPs contain more than 1 RBD. Non-canonical is abbreviated “Non-canon.” F. Bar plot analysis of the peptide peak areas derived from the MS data for csRBPs (orange) and all membrane proteins (gray) found across the four cell lines in (A). The peak area for each protein is individually plotted. The number of proteins in each set are annotated as numbers in the middle of each bar. Statistical differences between the RBPs and all membrane proteins was assessed with a Mann-Whitney test and p values are displayed on the plot. G. Flow cytometry analysis of K562 and OCI-AML3 (AML3) stained live with AF647-conjugated antibodies against the targets listed. Quantitative analysis of binding events was facilitated by company intensity to predefined AF647-conjugated beads and value converted into binding events based on the number of dyes per antibody. Binding events calculated from isotype antibody binding were subtracted from target values and plotted. Experiments were performed in biological triplicate.

To examine the distribution of proteins across cell types, we intersected the high confidence Mem-IP hits (**Figure 2C**). We found hundreds of unique proteins (except for A549) identified in the Mem-IP for each of the four cell lines (**Figure 2C**). To identify commonly presented csRBPs, we required an RBP to be present in 3 of 4 of our Mem-IP lists and found over 50.2 % (143 hits) of these proteins are RBPs (**Table S2**); even considering the most conserved list of proteins, requiring hits identification in all four cell lines, 57 of 98 (58%) of these proteins are RBPs (**Figure 2C**). We picked at least 3 cell lines as our criteria because the set of 4 cell lines span 2 suspension and 2 adherent. These criteria allows for some cell-type specific variation, as expected. There also exist 256 RBPs that were only identified in one of the cell lines analyzed, and 199 of these were associated with HEK293 cells. Finally, to examine if csRBP hits were more likely to be found as cell type specific (e.g. “293 unique”) or found in multiple cell surface proteomes (e.g. HEK293+ >=1 other) we calculated the fraction of RBPs singular vs. combined Mem-IP datasets. In each of the four cell lines, csRBPs were fractionally more represented across multiple cell lines compared to uniquely identified in a single cell line (**Figure 2D**). Together, these data indicate that many similar csRBPs appear on diverse cell lines.

To better understand the characteristics of csRBPs relative to the total pool of RBPs, we leveraged previously annotated RNA-binding domains (RBDs) to classify RBPs from the all-RBP list, the csRBP candidate list from the data-driven search, and RBPs identified in our cell-surface proteomics (“Mem IP RBPs”). Both csRBP lists exhibited a greater fraction of RBPs with non-canonical RBDs than the overall pool of RBPs, while being somewhat depleted for RBPs with RRM domains (**Figure 2E**). Moreover, it seems the set of putative csRBPs obtained from public databases (**Figure 1**) contains roughly three times as many ribosomal RBPs as those identified with our stringent cell-surface proteomics method (**Figure 2E**). Given that ribosomal proteins stand as some of the more highly expressed proteins in the cell and often appear as background in mass spectrometry experiments, the depletion of ribosomal proteins in our cell-surface proteomics suggests greater stringency and is consistent with high fidelity of the csRBP list generated from these experiments. A compiled list of high confidence csRBPs resulted in a total of 179 proteins and is detailed in **Table S2**.

We next asked how the apparent abundances of csRBPs compare to those of other cell surface proteins. Using the peak area detected by the mass spectrometer as a proxy for protein abundance (see **Methods**), we compared the average of peak areas of either csRBPs or all proteins identified in the Mem IP datasets, normalized for the amino acid length of each protein. This analysis showed that the csRBPs exhibit an average abundance similar to the total collection of cell surface proteins from each cell line (**Figure 2F**). The robust detection of peptides from RBPs in the Mem IP relative to the total pool peptides indicates that csRBPs may be significant components of the cell surface. However, differences in ionization and post-translational modifications could impact the relative detection across specific proteins.

To orthogonally assess cell-surface abundance of csRBPs and canonical membrane proteins, we employed a bead-calibrated live cell flow cytometry method. Canonical cell-surface proteins and csRBPs identified in our cell-surface proteomics were chosen as targets for immunolabeling and quantification. Live OCI-AML3 and K562 cells were stained with primary conjugated Alexa Fluor 647 (AF647) antibodies (NCL, hnRNP-A1, hnRNP-U, DDX21, DNA-PKcs, PRDX1, TFRC, STOM, LAT1, GLUT1, EZR, ꞵ2M, GYPA) and analyzed by flow cytometry. Quantification was achieved through the generation of a standard curve using beads bound with defined amounts of AF647 (see **Methods**). All csRBPs were detected at greater intensities than isotype controls in at least one cell line tested (**Figure 2G**). With the exception of TFRC, antibody binding between these csRBPs and the canonical membrane proteins tested remained within the same order of magnitude. GYPA, a late-stage erythroid cell surface marker, was highly expressed and largely restricted to K562 cells (**Figure 2G**), consistent with their erythroid state. ꞵ2 microglobulin (ꞵ2M), a major component of the MHC-I peptide presentation complex, was also highly abundant and poorly expressed on K562 cells (**Figure 2G**) as previously reported (Garson et al., 1985), confirming the specificity of this assay.

### Cell surface RNA binding proteins form distinct nanometer scale clusters

Like other cell surface protein (Fischer et al., 2021; Owen et al., 2010; Werbin et al., 2017), csRBPs were found to form clusters on the cell surface using diffraction-limited (DL) confocal imaging in Figure 1. To better understand the nanoscale organization of these clusters, we performed super-resolution (“SR”) single-molecule localization microscopy (**Figure S3A**). Using adherent cells to facilitate analysis, we collected data using primary conjugated anti-hnRNP-U-AF647 or anti-DDX21-AF647 antibodies applied to PANC1 or A549 cell lines (**Figure 3A-C, S3B**). Widefield fluorescence imaging demonstrated signal accumulating around the periphery of the cells consistent with our previous DL data. Using the imaging, post-processing, and data analysis steps outlined in Figure S3A (detailed in **Methods**) we generated SR reconstructions of three PANC1 cells (hnRNP-U), three A549 (hnRNP-U), and four A549 (DDX21) stained cells (**Figure 3A-C, and Table S3**). Examining the SR reconstructions from each of the three conditions revealed that the csRBPs formed distinct clusters of apparently uniform size. To measure the features of the SR reconstructions we implemented a series of quantitative analyses to define the precise features of the observed clusters. First, we implemented a semi-automated analysis workflow, performing Ripley’s analysis to determine the most common cluster size, which was then used as input for subsequent DBSCAN analysis (Nieves et al., 2023). The diameter of cs-hnRNP-U clusters were 139nm and 164nm on PANC1s and A549s, respectively, while csDDX21 clusters exhibited an average diameter of 133nm on A549 cells (**Figure 3D**). The average distance between clusters was most similar between hnRNP-U on PANC1s and DDX21 on A549s at ∼230 nm, while hnRNP-U on A549s had clusters that were slightly further apart at ∼316 nm (**Figure 3E**). Finally, we calculated the localizations per cluster, a measure of the number of dyes localized. We found that cs-hnRNP-U had an average of 36 dyes per PANC1 clusters and 61 dyes per A549 clusters, while csDDX21 had fewer, at an average of 12 dyes per A549 cluster (**Figure 3F**). The number of dyes localized is not a direct measure of ligand amount, but is correlated, suggesting more hnRNP-U antibody binding events on the cell surface as compared to the DDX21 antibody. While there are quantitative differences observed between these two cell types and two csRBPs, the overall cluster size is similar – within the diameter of 1 ribosome – raising the question of the relationship between these surface clusters and csRBP occupants.

**Figure 3.**
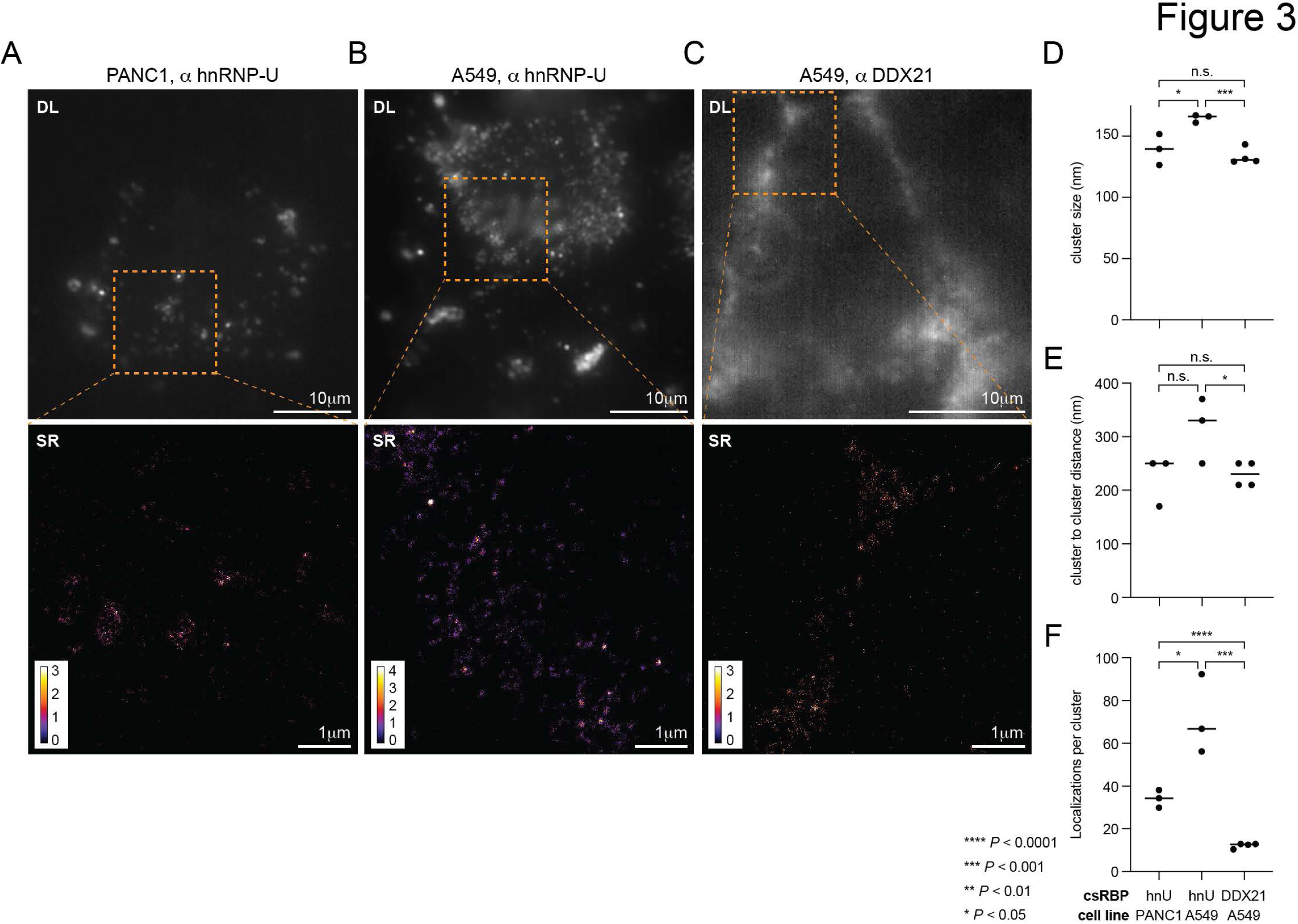
Super resolution reconstructions reveal clustered organization of csRBPs. A. PANC1 cells stained live with anti-hnRNP-U antibody, fixed, and then imaged with widefield microscopy at diffraction limited (DL, top) or super resolution reconstruction (SR, bottom). B. A549 cells stained and analyzed as in (A). C. A549 cells stained with anti-DDX21 antibody and analyzed as in (A). D. Bar plots of the quantification of the cluster size in nanometers (nm) for each cell. Median value is shown as a horizontal bar. Pairwise t-tests were performed between indicated datasets to evaluate the significance, p values are displayed. E. Bar plots of the quantification of the cluster to cluster distance in nm for each cell, plotted as in (D). F. Bar plots of the quantification of the points per cluster for each cell, plotted as in (D).

### csRBPs Cluster Together on the Cell Surface

Having found that the csRBPs form distinct and well-defined clusters on the cell surface using SR microscopy, we sought to understand what other protein factors assembled with the csRBPs. Using proximity labeling (Bar et al., 2018; Rees et al., 2015; Rhee et al., 2013) and adapting our previous method that enabled cell surface glycoRNA labeling (Flynn et al., 2021) we used biotin-phenol to tag cell surface proteins in proximity to various csRBP antibodies (**Figure 4A**). We selected the classically described csRBP of NCL as well as putative csRBPs DDX21 and PCBP1 for analysis (**Figure S4A, S4B**). We compared these csRBP profiles to the well-characterized and highly abundant cell surface protein ꞵ2M (**Figure S4C**). On OCI-AML3 there is anti-ꞵ2M labeling over isotype control with (**Figure S4C**) while this was not the case on K562 cells, consistent with reported poor expression of the MHC-I complex on K562s and our molecular counting data (**Figure 2G**). We also examined the specificity of this method by comparing the blotting signal after labeling K562 cells with an anti-DDX21 or anti-RIOK1, the latter we discovered was never identified as a csRBP (Figure 1). Performing proximity labeling on cells with the anti-RIOK1 antibody generated biotinylation signal similar to that of the isotype control (**Figure S4D**), additionally validating that most of our biotin signal was coming from intact cells and RBP antigens on the surface. After biotin enrichment, trypsin digestion, and MS analysis (**Methods**), we identified candidate molecular neighbors by quantifying the enrichment of each respective protein relative to proximity labeling with an isotype primary antibody (**Table S4**). From proximity labeling on OCI-AML3 and K562 cells, we identified GO terms consistent with membrane labeling and also many proteins with terms related to the cytosol, nucleus, and RNA binding (**Figure S4E**).

**Figure 4.**
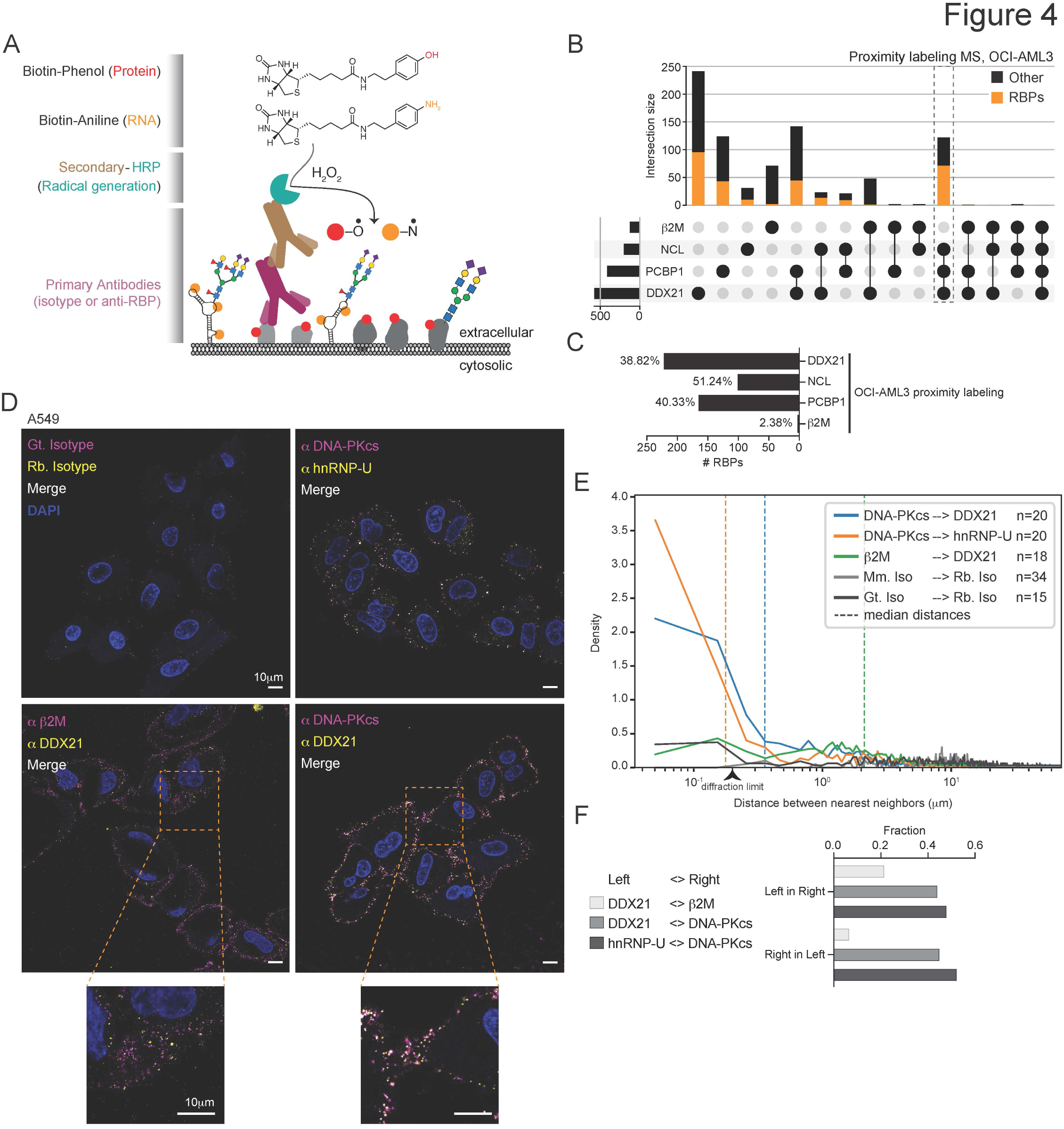
csRBPs co-assemble on the surface of living cells. A. Cartoon of HRP-based cell surface proximity labeling. Primary antibodies target specific cell surface antigens and recruit a secondary antibody conjugated to HRP. Upon addition of hydrogen peroxide, HRP converts biotin-aniline (for RNA labeling) or biotin-phenol (for protein labeling) into their radical form which is impermeable to intact cell membranes. Subsequently proteins or RNA can be analyzed for biotinylation. B. Intersection analysis visualized using an upset plot examining all hits identified from antibodies targeting anti-ꞵ2M, anti-NCL, anti-PCBP1, and anti-DDX21 on OCI-AML3 cells. The set size of each dataset is displayed as a bar plot at the left end of the intersection table. For each bar, the number of RBPs is overlaid in orange. A dashed box highlights the intersection of the three RBP datasets. C. Bar plot of the number (y-axis) and percent (annotated on each bar) of RBPs found in each of the proximity labeling datasets from (B) and (C). D. Confocal microscopy of A549 cells stained live and then fixed for analysis. Three color imaging was performed with target 1, target 2, and DAPI in purple, yellow, and blue, respectively. Overlapping signal of two target antibodies is displayed in white. Images were acquired using a 63x oil objective, a single z-slice is shown, and the scale bars are 10 µm. Zoomed regions are also marked with 10 µm scale bars. E. Nearest neighbor distance analysis of the antibody pairs imaged in (E). For each pair, the distance (nanometers, nm) from that anchor (left side protein name in the figure key) to the other pair was calculated across the indicated number of cells. These values were plotted in a density histogram and the mean distance is annotated with a dashed line. F. Bar plot analysis of a Manders coefficient calculation for the pairs imaged in (E). The fractional overlap of each pair, in both directions, were calculated and plotted.

To understand how data compared across surface protein targets, we intersected the molecular neighborhoods of each target antibody (**Figure 4B, S4F**). On OCI-AML3, the number of shared proteins between the three csRBP molecular neighborhoods (highlighted column) is much larger than the sum of common proteins between the ꞵ2M molecular neighborhood and those of csRBPs (**Figure 4B**). K562 data demonstrated a similar population, but comparison to ꞵ2M was not possible (**Figure S4F**). Further examining the proximity hits of OCI-AML3, we found an enrichment in RBPs in the molecular neighborhoods of csRBP anchors (38-51% of neighboring proteins). Conversely, approximately 2% of neighboring proteins of ꞵ2M were RBPs (**Figure 4D**). The lack of overlapping hits between ꞵ2M and the csRBPs suggests that csRBPs occupy distinct physical spaces from protein complexes like the MHC.

To test this hypothesis at the single-cell level, we employed confocal microscopy. We set up pairwise colocalization experiments with csRBP hits including DDX21, hnRNP-U, and DNA-PKcs as well as ꞵ2M as a control. To enable more robust imaging, we focused on the adherent A549 cell line for analysis. As seen in part in Figure 1 as well as in Figure 3 at much higher resolution, all of the csRBPs formed distinct puncta on the cell surface (**Figure 4D, S4G**). Using an object-based strategy to detect binding events, the pairwise analysis of the csRBPs to each other and ꞵ2M revealed strong colocalization between the csRBPs and much weaker overlap with ꞵ2M (**Figure 4D, 4E, 4F**). Most of the csRBP objects were within the diffraction limit of the confocal while more of the ꞵ2M pairs were not (**Figure 4E**, carrot at 0.2 µm denotes diffraction limit of light). Further examination of the cells stained with matched isotype antibodies does reveal some cell surface staining; however, it is weaker than csRBP or ꞵ2M staining (**Figure 4D**). To confirm these background spots did not follow similar colocalization patterns as seen for the csRBPs, we performed the same nearest neighbor analysis and found neither pairs of isotype antibodies localized near each other (**Figure 4E**). We next calculated the fractional overlap of each antibody cluster that colocalized with the other pair and found that approximately 45% of csRBP antibody spots were shared (e.g. 44% of all csDDX21 clusters were colocalized with csDNA-PKcs and 45% of all csDNA-PKcs clusters colocalized with csDDX21). However the fractional overlap was much less with ꞵ2M: 21% of csDDX21 clusters colocalized with ꞵ2M while only 6.8% of ꞵ2M clusters colocalized with csDDX21 (**Figure 4F**), suggesting that while some csDDX21 is near ꞵ2M, the vast majority of ꞵ2M exists in distinct domains on the cell surface. Taken together, these results suggest that RBPs cluster together on the cell surface.

### Cell surface RNA binding proteins colocalize with RNA, depend on RNA for clustering, and are in proximity to glycoRNAs

An accumulation of specific RBPs on the cell surface in distinct clusters raises the possibility that these domains may also contain cell surface glycoRNA, a logical molecular partner for an RBP. To address this possibility, we took three approaches. First, we assessed the colocalization of csRBPs and RNAs with confocal microscopy. We visualized both csRBPs by immunolabeling DDX21, hnRNP-U, and DNA-PKcs as well as dsRNA with the anti-dsRNA antibody 9D5 on live A549 cells (**Figure 5A**). Like other dsRNA-specific antibodies, 9D5 was initially described to bind RNA virus genomes (Kitagawa et al., 1977; Son et al., 2015) and like the J2 antibody, we predicted that if used on live cells, it could also detect the dsRNA regions of cell surface glycoRNAs. The 9D5 signal was organized in puncta around the periphery of the cell much like that of csRBPs (**Figure S5A**), revealing that this dsRNA antibody can detect RNA on the surface and it is also clustered. Examining the co-stains with the three csRBPs showed many of the 9D5 puncta overlapped with csRBP puncta (**Figure 5A, and insets**). We next applied object-based detection to quantify colocalization using nearest neighbors analysis, which demonstrated that many csRNA puncta lie within the optical diffraction limit of spots associated with all three csRBPs tested (**Figure 5B**). This result was similar to what was observed with csRBP imaging (**Figure 4**). By quantification, between 20-40% of all 9D5 clusters overlapped with clusters of csDDX21, cs-hnRNP-U, or csDNA-PKcs (**Figure 5C**), while the csRBPs clusters had approximately 8 to 23% overlap with 9D5 clusters (**Figure 5C**). Because anti-dsRNA antibodies like 9D5 have some differential specificity (Störk et al., 2021), these overlap data suggest that 9D5 can detect only a subset of the available csRBP clusters, but that many of the clusters 9D5 targets also contain csRBPs.

**Figure 5.**
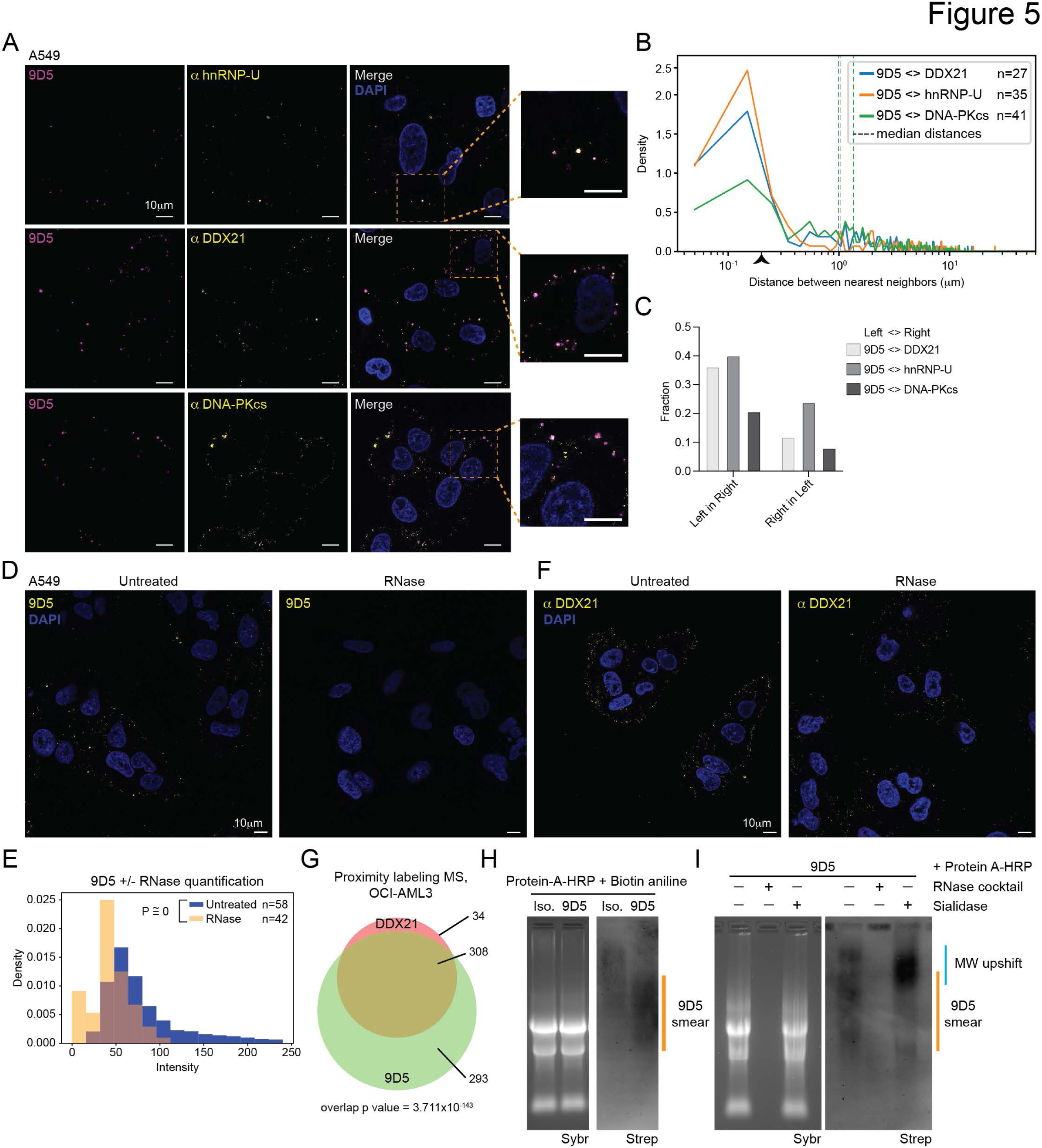
RBPs colocalize with and are dependent on glycoRNA on surface of living cells. A. Confocal microscopy of A549 cells stained live and then fixed for analysis. Three color imaging was performed with target 1, target 2, and DAPI in purple, yellow, and blue, respectively. Overlapping signal of two target antibodies is displayed in white. A single z-slice is shown and scale bars are 10 µm. B. Nearest neighbor distance analysis of the antibody pairs imaged in (A). For each pair, the distance (nm) from that anchor (left side protein name in the figure key) to the other pair was calculated across the indicated number of cells. These values were plotted in a density histogram and the mean distance is annotated with a dashed line. C. Bar plot analysis of a Manders coefficient calculation for the pairs imaged in (B). The fractional overlap of each pair, in both directions, were calculated and plotted. D. Confocal microscopy of A549 cells with and without RNase treatment and then stained live with 9D5 and then fixed for analysis. A single z-slice is shown, scale bars are 10 µm. E. Comparison of the distributions of 9D5 spot fluorescence intensity on A549s treated with or without RNase. The number of cells (n) quantified is shown for each condition. A p-value was calculated by assessing the statistical significance of the difference between the medians of the intensity distributions by using bootstrapping to perform a nonparametric permutation test. F. Confocal microscopy as in D, here staining with anti-DDX21 Scale bars are 10 µm. G. Venn diagram overlap of the proximity labeling hits identified using either DDX21 or 9D5 as antibody anchors on the surface of OCI-AML3 cells. The numbers labeled on the right indicate the set size for each overlapping region and the Mann-Whitney calculated p value for the overlap is shown. H. Cell surface proximity labeling assisted by Protein A-HRP and biotin-aniline to label cell surface RNAs. Isotype and 9D5 antibodies were used as anchors and the resulting total RNA (Sybrgold signal, Sybr) was analyzed on a northern blot, detecting biotinylated species (Streptavidin IR800, Strep). A 9D5-specific smear is highlighted in orange. I. In vitro digestion with RNase cocktail or sialidase of RNA isolated from 9D5-proximity labeling as in (H).

To establish the specificity of 9D5 for targeting cell surface RNAs, we performed the staining of live cells with and without pre-treatment with RNase A (single stranded RNase) and RNase III (double stranded RNase), which we predict would robustly remove cell surface RNA molecules. Live cell RNase treatment led to a > 90% reduction in 9D5 clustering on the cell surface (**Figure 5D, left**), as measured by total intensity per spot per cell (**Figure 5E**) and number of 9D5 spots per cell (**Figure S5B**). Because 9D5 binds near csRBPs, we also assessed the clustering of csDDX21 upon treatment of live cells with the RNases. We observed a more modest effect on the clustering of csDDX21 – reducing the clustering intensity by ∼60% (**Figure 5E right, S5C**) while the number of csDDX21 clusters was not significantly impacted (**Figure S5D**). To further investigate the association between csDDX21 and 9D5, we performed cell-surface proximity labeling for 9D5 on AML3 cells (**Figure 4A**, **S4B**). Intersecting 9D5 molecular neighborhoods with those of the csDDX21 revealed a high degree of intersection between the two neighborhoods on both cell types, 90% of the csDDX21 proximal proteins were also identified in the 9D5 dataset (**Figure 5G, Table S4**). Moreover, 32.6% of the 9D5 molecular neighborhood comprises csRBPs. These data are consistent with the hypothesis that cell-surface dsRNA and csRBPs are associated with each other.

Finally, we examined the features of the RNAs in proximity to the 9D5 antibody. We again used the proximity labeling method (**Figure 4A**) but employed biotin aniline to preferentially label RNA rather than proteins (Zhou et al., 2019). We found that a broad biotinylated smear of signal is detected in total RNA recovered from labeled AML3 cells. This labeling differed in molecular weight and had stronger signal intensity compared to isotype control labeling (**Figure 5H**). To determine if this signal could be attributed to glycoRNA, we repeated the assay but instead digested the isolated RNA *in vitro* with an RNase cocktail or, separately, a broad spectrum sialidase. The biotin smear was lost (sensitive to) RNase treatment while sialidase treatment resulted in a consolidating shift in molecular weight but no loss in signal intensity (**Figure 5I**). We previously observed this sialidase-dependent up-shifted signal when labeling cell surface glycoRNAs with MAAII or WGA lectin anchors (Flynn et al., 2021). Together these data suggest the presence of cell surface glycoRNA-csRBP clusters.

### Cell binding and entry of TAT-eGFP is dependent on cell surface RNA and RNA binding

The organization of cell surface molecules and complexes can impact the binding, signaling, or internalization of extracellular molecules as they encounter a target cell. CPPs, of which TAT was the first discovered, have many proposed mechanisms of cell surface binding and entry (Guo et al., 2016; Langel, 2022). TAT is a 9 amino acid peptide in the class of cationic CPPs, and its entry mechanisms have been characterized as dependent on negatively charged biopolymers like heparan sulfate proteoglycans (HSPG, (Rusnati et al., 1999; Tyagi et al., 2001; Urbinati et al., 2009). Our evidence for glycoRNA-csRBP clusters provides a new, but yet untested cell surface domain of poly-anions. We reasoned that processes dependent on glycoRNA-csRBP clusters might be modulated as a consequence of RNase addition. We therefore set out to assess how various forms of TAT functionally interact with cells treated with and without RNase.

Before entering cells, CPPs must interact with the cell surface. We assessed binding of a recombinant TAT-WT-eGFP fusion (**Figure S6A**) by incubating the cells with the protein and then using an anti-GFP antibody (which will be impermeable to live cells) to detect cell surface TAT-WT-eGFP (**Figure 6A**). TAT-eGFP forms puncta on the surface of cells, and upon 9D5 co-staining, we observed strong colocalization (**Figure 6B, 6C**). A control eGFP protein weakly binds the cell surface and when compared to 9D5 co-staining, there is little colocalization (**Figure S6B**), suggesting a specific co-binding activity of TAT-WT-eGFP to sites where 9D5 binds. We also assessed the colocalization surface bound TAT-WT-eGFP with hnRNP-U and DDX21 and found overlapping signals on A549 cells (**Figure S6C**); we noted that the median distances between TAT and csRBPs is similar to that between csRBPs and 9D5 (**Figure 5B**) suggesting similar biophysical features. To assess the dependence of RNA on TAT-eGFP binding, we pretreated live cells with a pool of RNases and again monitored the ability for TAT-WT-eGFP or eGFP to bind the cell surface while on ice. RNase treatment reduced the ability for TAT-WT-eGFP to bind to cells by ∼30% as compared to non-RNase treated cells (**Figure 6D, 6E**).

**Figure 6.**
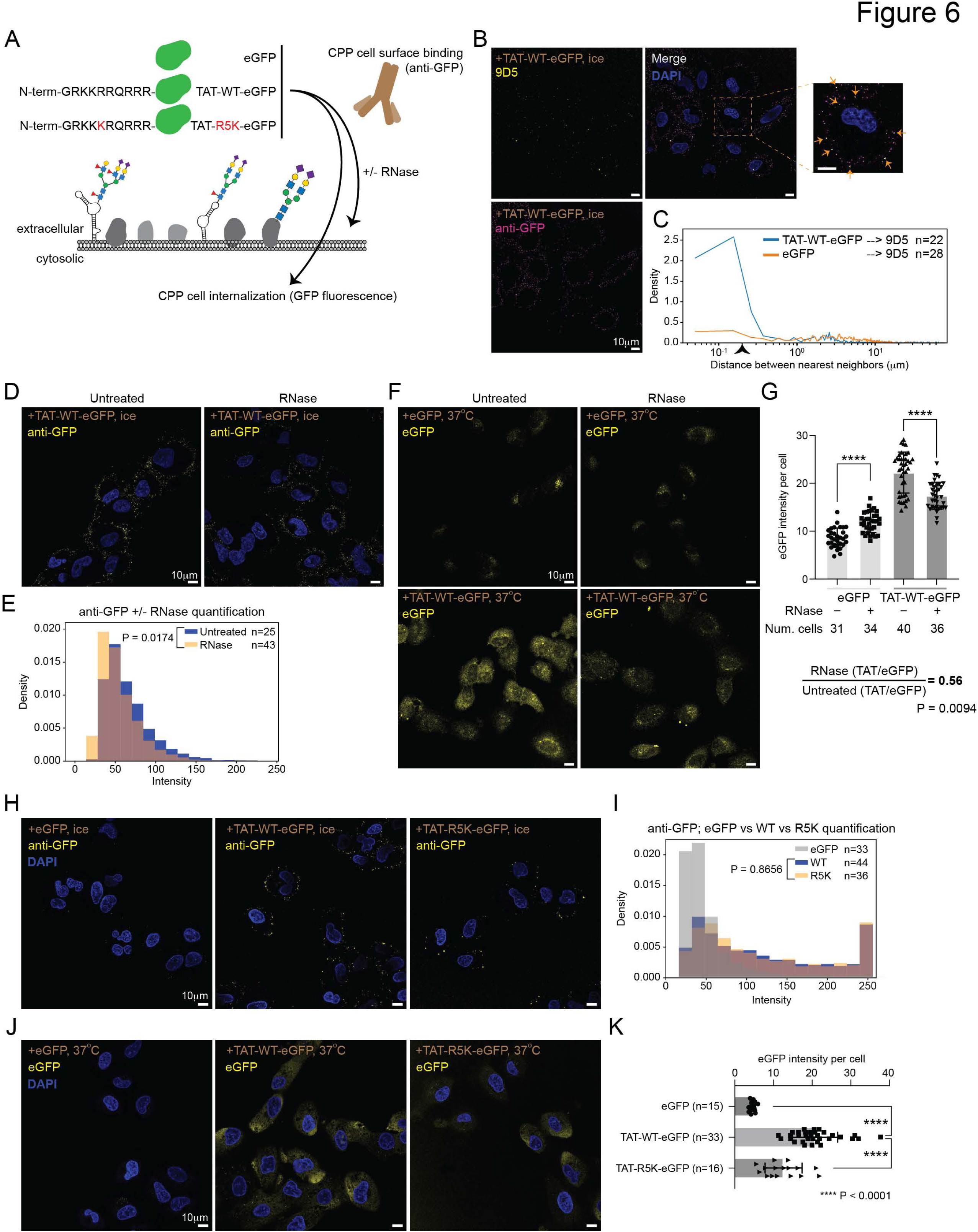
The RNA binding and cell penetrating TAT peptide is dependent on RNA for entry into cells. A. Cartoon of eGFP conjugate cell treatment and internalization assays. eGFP, TAT-WT-eGFP, or TAT-R5K-eGFP is added to cells with or without RNase. For cell binding, proteins are added while cells are on ice whereas for cell internalization the proteins are added to live cells in culture media at 37℃. B. Confocal microscopy of A549 cells with eGFP conjugates added on ice, then stained live with 9D5 (yellow) and anti-GFP (magenta) and then fixed for analysis. Overlapping signal is shown in white. The zoomed region is highlighted and overlapping 9D5-anti-GFP spots are noted with orange arrows. Scale bars are 10 µm. C. Nearest neighbor distance analysis of the antibody pairs imaged in (B). For each pair, the distance (µm) from that anchor (left side protein name in the figure key) to the other pair was calculated across the indicated number of cells and these values were plotted in a density histogram. D. Confocal microscopy of A549 cells as in B, here pretreated with or without RNase and stained only for analysis with anti-GFP (yellow). Nuclei are stained with DAPI (blue), scale bars are 10 µm. E. Comparison of the distributions of 9D5 spot fluorescence intensity on A549s treated with or without RNase. The number of cells (n) quantified is shown for each condition. A p value was calculated by assessing the statistical significance of the difference between the medians of the intensity distributions by using bootstrapping to perform a nonparametric permutation test. F. Confocal microscopy of A549 cells with eGFP conjugates added at 37℃, washed, fixed, and imaged for GFP fluorescence. Average projection images are shown and scale bars are 10 µm. G. Quantification of the per-cell GFP fluorescence intensity observed in (F). The number of cells quantified in each condition is shown and p values were calculated by a Mann Whitney test, ****, p < 0.0001. H. Confocal microscopy of A549 cells as in (B), here cells were treated with eGFP, TAT-WT-eGFP, or TAT-R5K-eGFP and stained only for analysis with anti-GFP (yellow). Nuclei are stained with DAPI (blue), scale bars are 10 µm. I. Analysis as in (E) of the cell binding spot intensities from (H). J. Confocal microscopy of A549 cells as in (F), here cells were treated with eGFP, TAT-WT-eGFP, or TAT-R5K-eGFP. Nuclei are stained with DAPI (blue), scale bars are 10 µm. K. Analysis as in (G) of the total internalized eGFP fluorescence intensity from (J).

Most work investigating CPPs focuses on their ability to enter cells, resulting in conjugated molecules gaining access to the intracellular space. We next assessed if pretreatment of cells with RNase could impact the ability for TAT-WT-eGFP to translocate into live cells in culture media. After adding RNase and removing it from the live cell cultures, TAT-WT-eGFP or eGFP were added to conditioned culture media and allowed to incubate with cells for 20 minutes in standard growth conditions. TAT-WT-eGFP robustly enters the cell, developing cytosolic and nucleolar GFP signal as expected while eGFP has less potent cell entry capabilities (**Figure 6F, 6G, S6D**). Examining the RNase-treated cells revealed a 20% loss of TAT-WT-eGFP internalization, compared to the non-RNase treated cells (**Figure 6F, 6G, S6D**). RNase treatment conversely caused a slight increase in eGFP internalization resulting in the TAT-WT-eGFP+RNase levels to be closer to baseline internalization of a non-CPP containing eGFP (**Figure 6G**). Normalizing for this effect demonstrates that live cell RNase treatment causes a 47.5% reduction in the internalization of TAT-WT-eGFP compared to untreated cells (**Figure 6G**). These data suggest that in addition to binding, cell internalization of TAT-WT-eGFP may be controlled in part by cell surface glycoRNA-RNP clusters. To address this possibility without the need for modification of the cells, we took advantage of a well-described feature of the TAT CPP. Specifically, TAT contains specific arginines that are critical for transactivation of the HIV1 genome (Calnan et al., 1991a, 1991b) and TAR RNA binding (Gotora et al., 2023). To explore the possibility that the RNA-binding drives TAT cell binding or internalization, we mutated R52 (the 5^th^ amino acid in our peptide) to a lysine (K, **Figure S6A**), which maintains the same net charge in the mutant peptide while mutating a residue critical for RNA binding. When we examined the relative ability for eGFP, TAT-WT-eGFP, or TAT-R5K-eGFP to bind the surface of cells, we found no defect with the R5K mutation compared to the WT TAT sequence (**Figure 6H, 6I**). However, our cell entry assay revealed that TAT-R5K-eGFP’s internalization was reduced by nearly 40% as compared to TAT-WT-eGFP (**Figure 6J, 6K**).

## Discussion

Here we explored the composition and organization of the mammalian cell surface in the context of glycoRNA biology. Employing several orthogonal molecular and imaging tools, we demonstrated that many RBPs are found clustered on the cell surface in proximity to glycoRNAs. We also find that exogenous peptides like the CPP TAT can bind glycoRNA-csRBP clusters and that TAT’s ability to bind and enter cells is dependent on the integrity of these glycoRNA-csRBP clusters. These findings provide evidence that clustered glycoRNAs and csRBPs functionally regulate the internalization of extracellular factors.

## The cell surface as a platform for RNA biology

We were motivated to understand what could anchor cell surface glycoRNAs, and in considering this question we systematically evaluated the presence of RNA binding proteins on the cell surface. Our interest in generating a careful inventory of csRBPs was further motivated by several individual reports of RBPs detectable at the cell surface. As noted, NCL was first discovered as a cell surface protein in 1990’s (Semenkovich et al., 1990), but there have been other reports across the years highlighting this phenomenon. In our initial work describing glycoRNAs, we highlighted the fact that the RNA transcripts modified with glycans were correlated with known RNA autoantigens (Flynn et al., 2021). This connection is further extended in the case of csRBPs as the Ku proteins have been found on the cell surface (Dalziel et al., 1992; Prabhakar et al., 1990). The concentration of glycoRNAs and csRBPs in the same domains on the cell surface raise the possibility that these are sites of interaction for immune cells that could participate in the initiation or maintenance of autoimmune pathologies. Better defining how these glycoRNA-csRBP clusters are formed and regulated will contextualize their role in cell biology.

By cataloging various published datasets (**Figure 1**) we found that the type of chemistry used to enrich surface proteins had an impact on the number of csRBPs identified. Of specific interest is the periodate-based methods which should have selectively for sialic acid labeling. In this context and using the conservative need to be found in 5 or more datasets, we found 187 csRBPs enriched in periodate datasets (**Figure 1**). This suggests that many csRBPs are glycoproteins and could play roles on the cell surface like other more well studied glycoprotein ligands in cell-cell communication. In support of our systemic analysis, previous studies have found both N– and O-glycopeptides from NCL (Carpentier et al., 2005) and other RBPs (Hofmann et al., 2015). More generally, the experimental framework we use can extend to other unexpected proteins detected in cell surfaceome MS datasets, outside of the RBP family of proteins. For example, cytosolic metabolic enzymes are reproducibly found in surfaceome datasets (**Tables S1 and S2**). Careful validation by orthogonal methodologies such as live-cell staining and analysis as well as chemical proteomics strategies can provide robust evidence. Extending this past a binary readout of detection or no-detection, we worked to understand the relative quantity of csRBPs as compared to other more classically defined cell surface proteins by performing various quantitative analyses (**Figure 2**). We demonstrated that by both the number of proteins and their abundance, RBPs make up a robust portion of the cell surface. Importantly, we can detect full-length proteins by Western blot, indicating that the csRBP signal we are evaluating is not coming from MHC-presented peptides. This is further supported by our colocalization analysis which revealed poor overlap between ꞵ2M and other glycoRNA-csRBP clusters.

The presentation of full length and possibly folded RBPs opens new hypotheses surrounding how and what RNAs they directly interact with as well as in what context csRBPs are presented. Much of the MS data we compiled from published data were from cell culture models (**Table S1**) as was our NHS-fraction based surfaceome datasets (**Figure 2, Table S2**). Beyond this, the efforts here were initially focused on csRBPs that were found in many datasets, but exploring cell-type or –state specific csRBP expression could open up a new understanding of how the cell communicates to the extracellular space. When considering nucleic acids outside of the cell, there is a growing body of literature characterizing extracellular, nonvesicular RNAs (exRNAs, (Chai et al., 2023; Hoy and Buck, 2012; Murillo et al., 2019)). Despite clear evidence for presence of exRNAs across a wide variety of biofluids, little is known about how or if they specifically interact with the cell surface. The broad scope of our observation of csRBPs raises the possibility that these cell surface proteins could serve as binding patterns for exRNAs. Future work will be needed to define this putative interaction.

### Spatial organization of csRBPs and glycoRNAs

The assembly of molecular complexes enables regulation and in the context of the cell surface this happens both in cis (on the cell of origin) as well as in trans (between two cells). As analytical tools have evolved, our understanding of the organization of cell surface proteins has dramatically improved. We examined the physical distribution of csRBPs across a number of cell lines and with both confocal and super resolution reconstructions. Our diffraction limited studies demonstrated hundreds of puncta of csRBPs that often co-localize with one another, as well as the anti-RNA antibody 9D5. While 9D5 does not have glycoRNA specificity per se, proximity labeling demonstrated that 9D5 is molecularly adjacent to glycoRNA species on the cell surface (**Figure 5**). Focusing on DDX21 and hnRNP-U, we went below the diffraction limit of light and obtained nanometer scale reconstructions of these csRBPs (**Figure 3**). What was only clear after quantitative analysis of these data were that the csRBPs form highly regular clusters: both in the size of clusters (120-165nm) and the distance between clusters (230-300nm). These quantitative features were similar across RBPs and cell lines and prototypical sizes and organization of clusters could suggest a biologically regulated process giving rise to the observed spatial organization of the csRBPs and glycoRNAs in these domains. Traditionally, it has been assumed that lipid-lipid interactions are the driving force in membrane organization at the molecular level. However recent evidence (Shelby et al., 2023) suggests that the biology is more complex, with proteins and lipids affecting each other and driving membrane organization. Our data suggests that there is another layer of complexity, with cell surface tethered glycoRNA and RBPs operating to control the organization of select nano-clustered domains as well.

Further, a recently reported method that uses proximity ligation between a sialic acid DNA aptamer and an antisense RNA probe to probe where specific glycoRNA transcripts (termed Aptamer and RNA in situ Hybridization-mediated Proximity Ligation Assay, ARPLA) occur at subcellular resolution found that glycoRNAs form clusters on the cell surface (Ma et al., 2023). This is consistent with our initially proposed model of glycoRNAs (Flynn et al., 2021) and is further supported by our data here. Our characterization of glycoRNA-csRBP clusters suggests that these are likely the same domains visualized with this new technology and that RBPs co-populate glycoRNA clusters seen. Combining these insights to understand which RBPs are paired with specific glycoRNA transcripts could be accomplished through a similar strategy.

### RNA-dependency of cell penetrating peptide binding and entry

The ability for CPPs loaded with cargo to enter cells has driven significant interest in discovering new CPPs, understanding their properties, and engineering them for broad spectrum use as tools and delivery modalities (Guo et al., 2016; Langel, 2022). A major dependency on heparan sulfate proteoglycans (HSPGs) has been established to regulate cationic CPPs on the basis of non-specific charge interactions. Interestingly, while most published mechanisms of TAT cell interactions suggest HSPGs as the main cell surface ligand, there is some evidence for non-HSPG TAT cell activity (Gump et al., 2010). Because of the charge similarity of RNA and HSPGs as well as the known intracellular RNA binding role of TAT, we explored how our newly characterized glycoRNA-csRBP clusters might impact CPP biology. Our data demonstrating the RNase dependency of glycoRNA-csRBP clusters, TAT cell surface binding, and TAT entry suggests that the structure or assembly of these glycoRNA-csRBP clusters is important for the mechanism of TAT’s interaction with and eventual entry into cells. Since TAT was the first described CPP and has an important role in the HIV1 life cycle, there are detailed mechanistic insights, down to the amino acid level, of how the TAT protein operates. We leveraged this, and in particular the knowledge surrounding a critical arginine residue (R52, (Calnan et al., 1991a, 1991b; Gotora et al., 2023)) to break its RNA binding while maintaining net positive charge of the peptide. With this mutant form, we found a selective defect in the cell internalization but not cell binding, suggesting that the ability to form robust and specific RNA-protein interactions drives the productive association with the cell surface for cell entry.

Beyond TAT, there are many CPPs, some engineered and others naturally occurring. Among those that naturally occur, examples such as C9orf72 and the prion protein (PrP) are interesting to consider. C9orf72 can contain pathogenic G4C2 repeats that when translated result in dipeptide-repeats including poly-PR and GR, which have been shown to cause neuronal cell death in vitro and in vivo (Kwon et al., 2014; Mizielinska et al., 2014). Upon cell entry poly-arginine motifs can interact with RNA, localize to the nucleolus, and disrupt translation (Kanekura et al., 2018). In contrast, in the context of neurotoxic Amyloid-β peptide (negatively charged N-terminus with a hydrophobic C-terminus) aggregation, a signal-peptide derived CPP can act as an anti-prion agent (Löfgren et al., 2008; Söderberg et al., 2014). Here, the anti-aggregation properties are ascribed to the hydrophobic domain and cell entry to the cationic domain. The initially described cationic domain was KKRPKP and contains an arginine that could facilitate RNA binding, however a different motif KKLVFF had similar anti-amyloid effects (Henning-Knechtel et al., 2020). These examples highlight other biological contexts where the RNA-binding feature of cationic CPPs may regulate their cell association and entry, and could provide new ways to address neuroprotection or neurotoxicity by considering the role for cell surface glycoRNA-csRBP clusters.

Taken together we propose the idea of a new mechanistic dependency for cell entry: RNA-dependent import system (RIS). This pathway leverages these glycoRNA-csRBP clusters, defined here as sites of initial cell binding, and likely concentrates positively charged proteins or peptides like TAT into these domains. After cell binding, various mechanisms of internalization would be possible as the captured molecules could be brought close enough to the membrane for direct internalization or be taken up through endocytosis. The ability for RNase to destroy the clusters of glycoRNA-csRBPs as well as reduce the entry of TAT, suggests that their formation is somehow important for either mechanism of entry. Recently, it was shown that Cas9 RNPs with 4 copies of the SV40 NLS (PKKKRKVEDPYC) which harbor an arginine in position 5 (similar to TAT), enabled editing of mouse neurons in vivo with low adaptive immune response (Stahl et al., 2023). Better understanding the composition of the glycoRNA-RBP clusters, their formation, molecular architecture, and genetic dependencies will more completely define these new cell surface domains and could have impacts on functional cell delivery.

## Limitations

While we attempt to look broadly at what proteins are presented on the cell surface, we are limited to the reactivity of the chemical reagents used. Most of the data we analyzed leverages NHS (lysine), periodate oxidation of diols (glycans and mostly sialic acid), or radicalized phenol (tyrosine) labeling. As such, proteins that are poorly sialylated or have few surface lysines or tyrosines will not be efficiently sampled. We therefore suspect our data and analysis should serve more as a baseline rather than the full scope of what is presented on the cell surface. However, we do have confidence in comparing hits across conditions using the same labeling technology (e.g. looking at proteins in proximity to different RBP antibody anchors while using biotin-phenol). Expanding accessible chemistries and performing multiple labelings on the same cell or tissue models to better assess the composition will be important for future studies. Beyond this, the quality of the peptides, ability to analyze glycopeptides, and the type of MS used can greatly impact the number of high confidence peptide spectral matches which can modulate the depth to which any chemical labeling strategy can reveal a complete surface proteome.

The RNA binding properties of csRBPs lead us to explore the hypothesis of the importance of RNA binding in the interaction and internalization of the TAT, a cationic CPP. While we characterized a dependency of TAT’s cell surface binding and evential internalization on glycoRNA-csRBP clusters, we have not explored the relationship between other CPP classes (hydrophobic or amphipathic) and glycoRNA-csRBP clusters. Given that some amphipathic CPPs contain polar amino acids it is possible this class may also colocalize with the glycoRNA-csRBP clusters on the cell surface, however future work will be needed to define these features.

## Figure Legend

**Figure S1.**
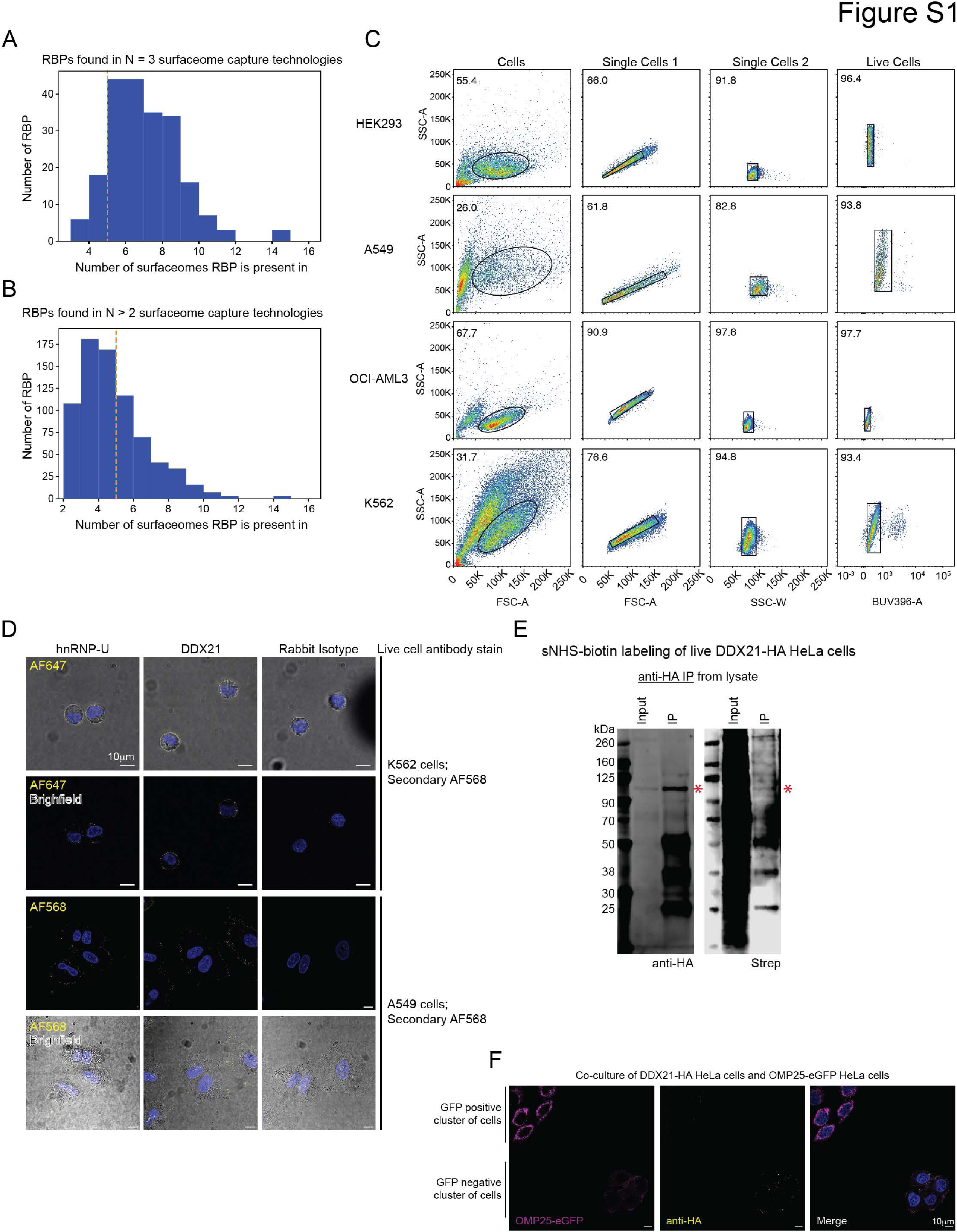
Validation strategies for establishing RBPs as cell surface proteins. A. Histogram plotting the number each RBP (y-axis) was found in a given set of cell surface proteomics experiments (surfaceomes, x-axis). RBPs were only considered for this analysis if they were identified in the three main classes of surfaceome profiling techniques (“N = 3”, **Methods**). B. Histogram plotting as in (A). Here RBPs were only considered for this analysis if they were identified in two of the three main classes of surfaceome profiling techniques (“N > 2”). C. Scatter plots of flow cytometry data. Cell lines are presented as rows and specific gates for cells, single cells1, single cell2, and live cells are presented as columns. The axes are labeled with the specific laser channels used for each analysis. Circles and boxes annotated within each scatter plot represent the population of cells sequentially analyzed and number in the top left indicates the percent of cells within that circle or box. D. Brightfield scans of the confocal images presented in Figure 1E. csRBP signal is overlaid to demonstrate the accumulation of csRBP signal at the periphery of the cells. E. Western blot analysis of HeLa cells expressing HA-tagged DDX21. Cells were labeled with sulfo-NHS-biotin before lysis, membrane organelles isolated, and HA-tagged proteins enriched. Input lysate and enriched material was subjected to anti-HA (left) or streptavidin (right) blotting. The expected molecular weight for DDX21 is noted with a red asterisk. F. Confocal microscopy images of immunolabeled HA-tag on live-stained DDX21-HA HeLa cells and OMP25-eGFP HeLa cells grown in coculture. Cells were plated in a 1:1 ratio and grown for 48 hours before being immunolabeled live on ice with anti-HA antibody and visualized with secondary antibody conjugated with AlexaFluor 568. OMP25-eGFP, HA-tag, and DAPI are displayed in magenta, yellow, and blue respectively. Images were acquired using a 63x oil objective, a single z-slice is shown, and the scale bars are 10 µm.

**Figure S2.**
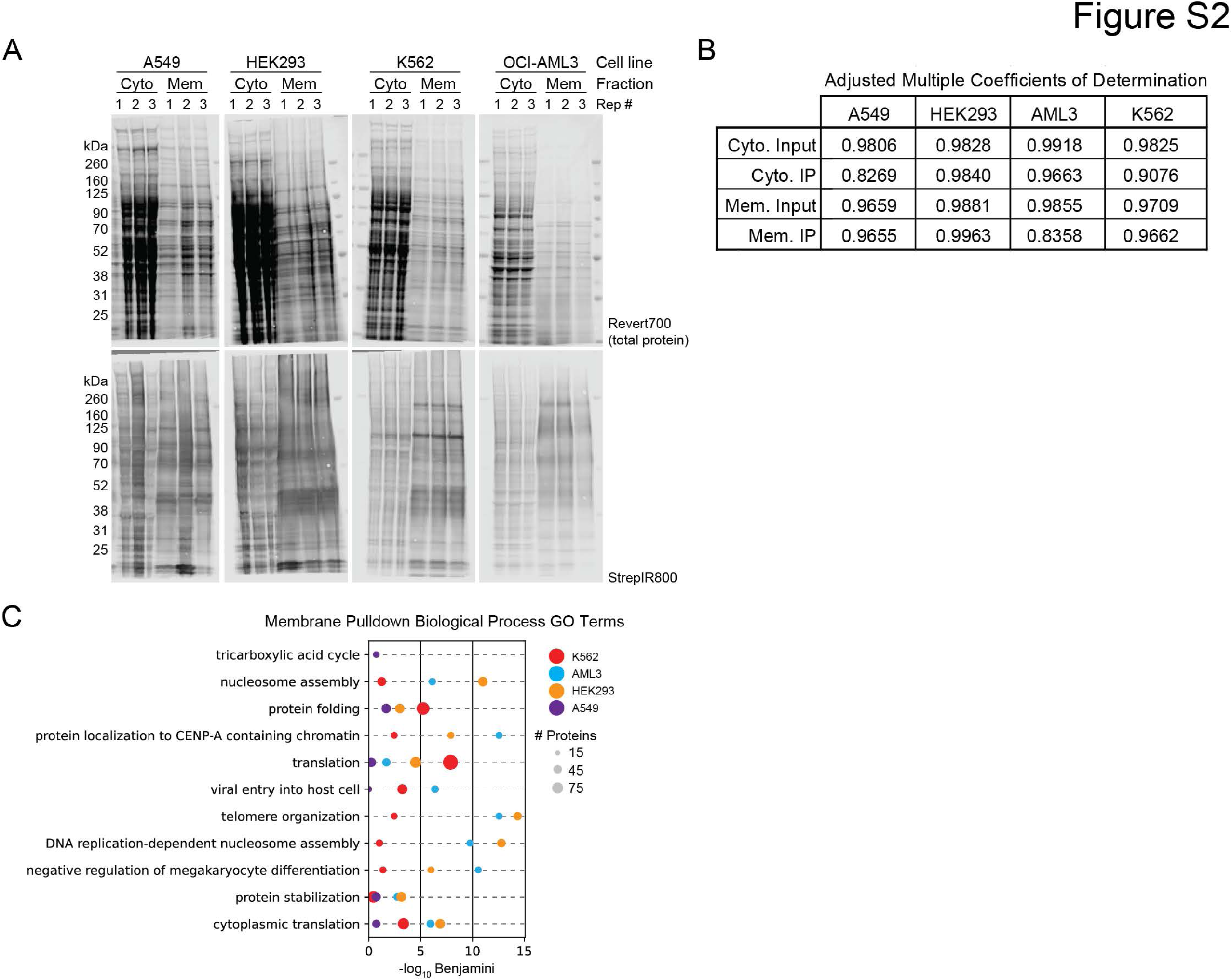
Biochemical fractionation enables deep cell surface proteomics. A. Western blot analysis of cytosolic (Cyto) and membrane (Mem) protein lysates isolated from A599, HEK293, K562, and OCI-AML3 cells after treatment with sulfo-NHS-SS-biotin before lysis. Biological triplicates are shown numbered 1, 2, and 3. Total protein signal is displayed in the upper row and biotinylated proteins visualized with Streptavidin-IR800 (Strep) in the bottom row. B. Table of the correlation analysis of the mass spectrometry input and enriched samples generated from the Cyto and Mem fractions from the four cell lines in (A). An adjusted multiple coefficients of determination analysis was performed to examine the pairwise similarity across the biological triplicates. C. Gene ontology (GO) biological process (BP) analysis of membrane enriched (Mem-IP) proteins identified by MS from HEK293, A549, K562, and OCI-AML3 (AML3) cells. The top 5 BP terms across the four cell lines were intersected and the union is displayed with the number of proteins in each term for each cell line (colors) represented by circle size and plotted on the x-axis by their significance. D. Gene ontology (GO) molecular function (MF) analysis as in (C).

**Figure S3.**
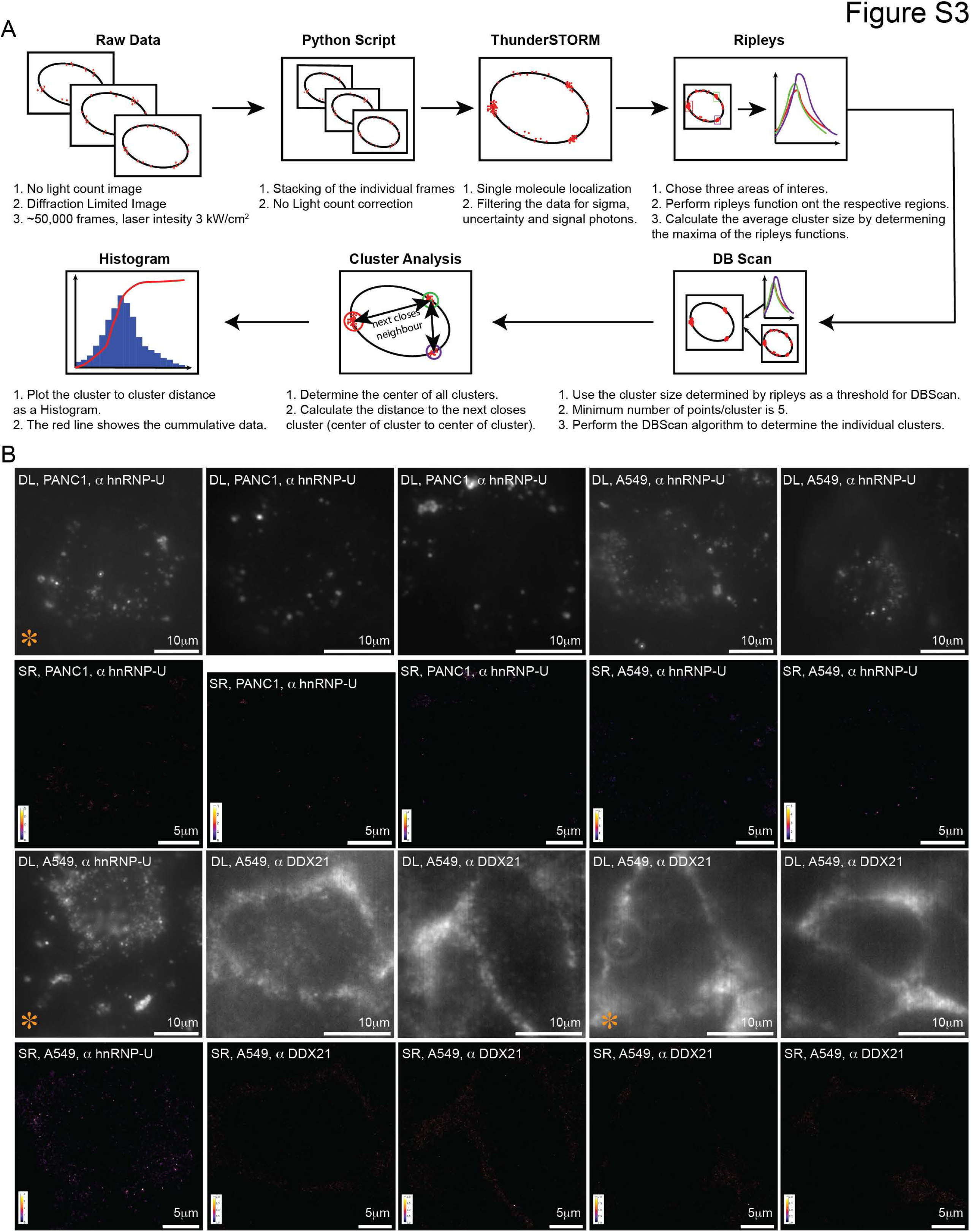
ThunderSTORM, Ripleys, and DBSCAN analysis enables super resolution reconstructions. A. Cartoon of the experimental, data acquisition, and data analysis pipeline used to collect and present DL and SR images. B. Individual DL (top rows, gray scale) and SR (bottom rows, colored scales) images of each individual cell collected for analysis. Scale bars are noted on each image. For SR panels, individual color bars are provided to indicate the relative signal intensity. Three cells have orange asterisk in the bottom left corners indicating these are the cells presented in Figure 3. The cell type and antibody stain are noted in the top left corner of each panel.

**Figure S4.**
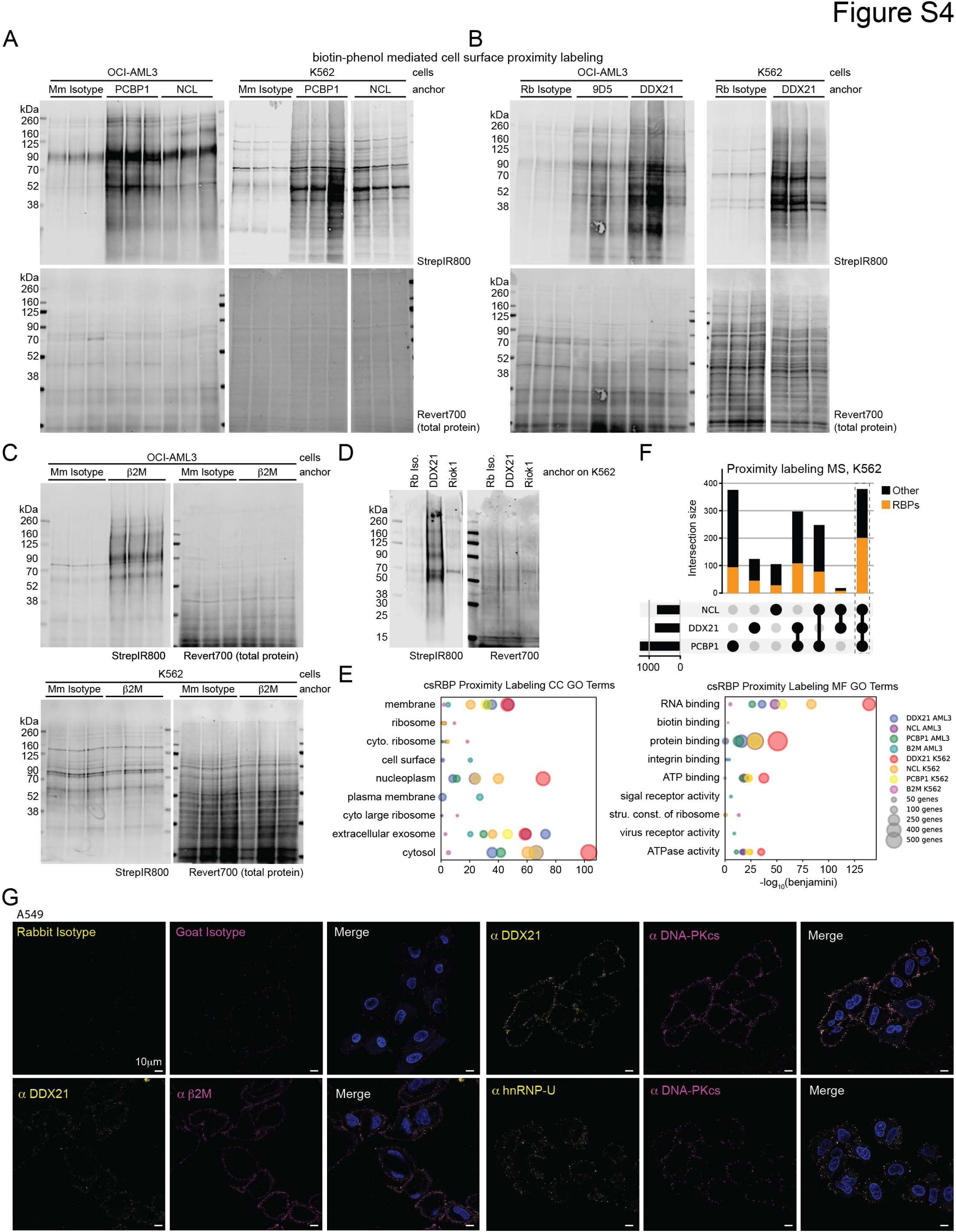
A. Western blot analysis of OCI-AML3 (left) and K562 (right) cells after cell surface biotinylation with biotin-phenol. Mouse host antibody anchors used in each condition are labeled and each lane represents an independent biological replicate. Biotinylated proteins are visualized with StrepIR800 and total protein loading assessed using Revert700. B. Western blot analysis as in (A), here showing rabbit host antibody anchors. C. Western blot analysis as in (A), here showing a second set of mouse host antibody anchors. D. Western blot analysis as in (A), here showing the Rabbit isotype, DDX21, and RIOK1 anchors, the last of which is an RBP not found as a csRBP in Table S1. E. Gene ontology (GO) cellular compartment (CC) and molecular function (MF) analysis of csRBP proximity labeling MS data from OCI-AML3 and K562 cells. The top 5 BP terms across the four anchor antibodies and two cell lines were intersected and the union is displayed with the number of proteins in each term for each cell line (colors) represented by circle size and plotted on the x-axis by their significance. F. Intersection analysis visualized using an upset plot examining all hits identified from antibodies targeting anti-NCL, anti-PCBP1, and anti-DDX21 on K562 cells. The set size of each dataset is displayed as a bar plot at the left end of the intersection table. For each bar, the number of RBPs is overlaid in orange. A dashed box highlights the intersection of the three RBP datasets. G. Confocal microscopy as shown in Figure 4D, here single-channel panels are shown. Three color imaging was performed with target 1, target 2, and DAPI in purple, yellow, and blue, respectively. Overlapping signal of two target antibodies is displayed in white. Images were acquired using a 63x oil objective and the scale bars are 10 µm.

**Figure S5.**
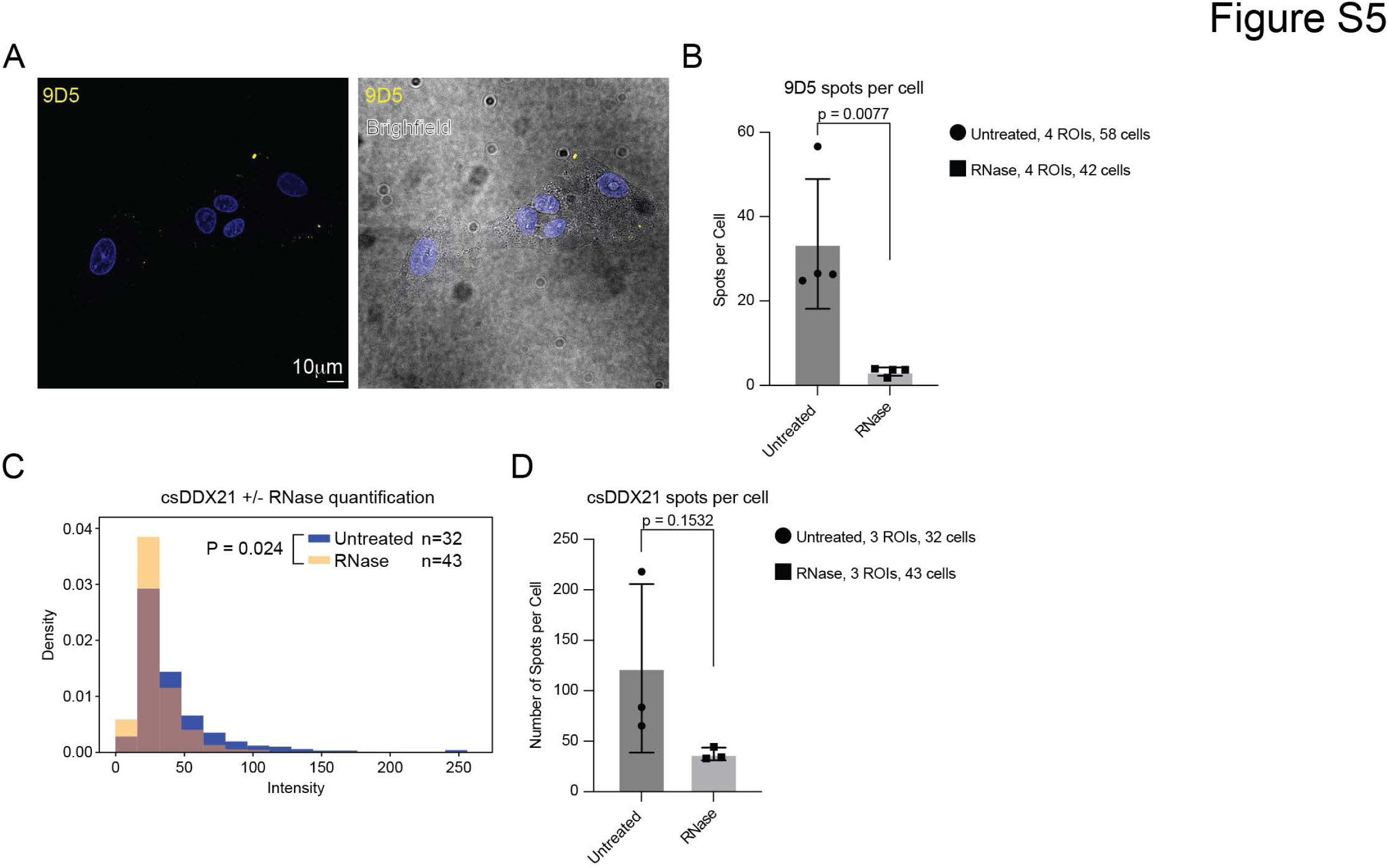
A. Confocal microscopy of A549 cells stained live and then fixed for analysis. Only the 9D5 antibody was used to stain these cells and then DAPI in blue and the bright field channels are overlaid to give cell surface context for the 9D5 clusters. Scale bar is 10 µm. B. Bar plot analysis of the number of 9D5 spots detected per cell on untreated or live cell RNase treated A549 cells. We quantified across 4 independent imaging regions (regions of interest, ROIs) aggregating the total number of cells indicated for each condition. Standard deviation is shown and an unpaired t test was used to assess statistical significance, p value shown. C. Comparison of the distributions of anti-DDX21 spot fluorescence intensity on A549s treated with or without RNase. The number of cells (n) quantified is shown for each condition. A p value was calculated by assessing the statistical significance of the difference between the medians of the intensity distributions by using bootstrapping to perform a nonparametric permutation test. D. Bar plot analysis as in (B), here quantifying the number of csDDX21 spots We quantified across 3 independent ROIs. Standard deviation is shown and an unpaired t test was used to assess statistical significance, p value shown.

**Figure S6.**
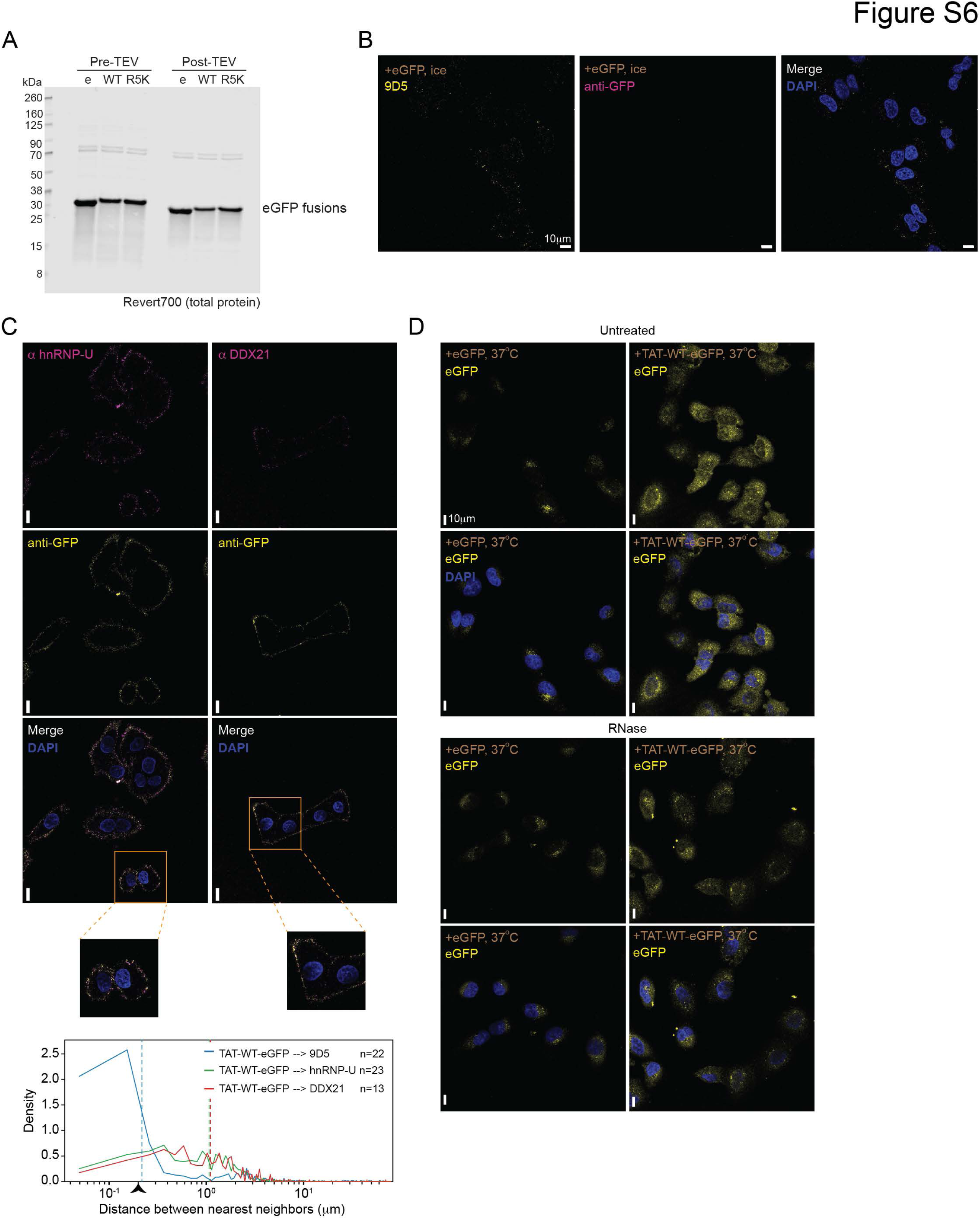
Expression and characterization of eGFP fusion proteins. A. Western blot analysis of expressed proteins eGFP (e), TAT-WT-eGFP (WT), and TAT-R5K-eGFP (R5K) after HIS-tag capture, before and after TEV protease cleavage. Total protein levels assessed using Revert700. B. Confocal microscopy of A549 cells stained live with eGFP fusion proteins (indicated in brown text) for 20 minutes on ice, washed, and then stained live with anti-GFP (magenta) or 9D5 (yellow), stained with DAPI (blue) and fixed for analysis. Overlapping signals of anti-GFP and 9D5 appear white. Scale bar is 10 µm. C. Confocal microscopy of A549 cells with eGFP conjugates added on ice, then stained live with anti-hnRNP-U (left, magenta) or anti-DDX21 antibody (right, magenta) and anti-eGFP (yellow) and then fixed for analysis. Overlapping signal is shown in white. The zoomed region is highlighted, and scale bars are 10 µm. Nearest neighbor distance analysis of the antibody pairs imaged in (C). For each pair, the distance (µm) from that anchor (left side protein name in the figure key) to the other pair was calculated across the indicated number of cells, and these values were plotted in a density plot. D. Confocal microscopy of A549 cells treated live with eGFP fusion proteins (indicated in brown text) for 20 minutes in culture media at 37℃, washed, fixed and stained with DAPI (blue) for analysis. eGFP fluorescence (no antibody) is shown in yellow. Pairs of images are shown with and without DAPI overlay.

## Table Legend

**Table S1.** RBPome and cell-surface proteome analysis.

**Table S2.** Cell-surface proteomics with sulfo-NHS-SS-biotin and subcellular fractionation.

**Table S3.** dSTORM image settings and quantitation.

**Table S4.** Cell-surface proximity labeling proteomics.

**Table S5.** Plasmid sequences for cell-free expression of eGFP and TAT-eGFP fusion proteins.

## Methods

### Cell culture

All cells were cultured 5% CO_2_ and 37℃. Adherent cell lines were maintained by first rinsing cells with 1x PBS and then treating cells with TrypLE (ThermoFisher Scientific) for a maximum of 5 minutes at 37℃ before quenching the reaction with complete media. Adherent cell cultures were split when cell density reached 80-90%. Adherent cells used here included HeLa (ATCC), HeLa-DDX21-HA (Calo et al., 2015), A549 (ATCC), and HEK293 (ATCC) and were all grown in 1x DMEM based media (ThermoFisher Scientific) with 1% Penicillin and Streptomycin (Pen/Strep, ThermoFisher Scientific) and 10% Heat Inactivated Fetal Bovine Serum (FBS, ThermoFisher Scientific). Adherent cells used for SR imaging were cultured in 1x RPMI-1640 based media (ThermoFisher Scientific) supplemented with 10% FBS (not heat inactivated, ThermoFisher Scientific), 1% GlutaMAX (ThermoFisher Scientific), and 1% Penicillin-streptomycin antibiotic cocktail (Sigma Aldrich). The Suspension cell lines were maintained by spinning cells down, removing the medium, and resuspending cells in fresh complete media. Suspension cell cultures were split when cell density reached 2 million cells per mL. Suspension cells used here included OCI-AML3 (obtained from the Sanger Institute Cancer Cell Collection) and K562 (ATCC) and were grown in 1x RPMI-1640 base media (ThermoFisher Scientific) with 1% Pen/Strep and 10% FBS. Cell cultures were periodically checked for and maintained as mycoplasma negative.

### DDX21-HA HeLa Cell and OMP25-eGFP Cell HeLa Cell Coculture

HeLa cells expressing DDX21-HA or OMP25-eGFP (expressed using pMXs-3XHA-EGFP-OMP25, Addgene: plasmid #83356) were cultured in 1x DMEM based media (ThermoFisher Scientific) as described above. These cells were trypsinized in TrypLE (ThermoFisher Scientific) for 5 min, resuspended in the media used for their culture, and counted. 5000 cells from each line were plated together, in a 8-chambered coverglass system (Cellvis, C8-1.5H-N), treated with 500 µL fresh media, and mixed by pipetting up and down five times, and incubated for 48 hours at 37℃. These cells were then immunolabeled with rabbit anti-HA antibody (Abcam, ab9110) live on ice, fixed, treated with DAPI, and imaged as described below.

### Flow cytometry and cell staining

Cells were cultured as above and directly counted (suspension cells) or gently lifted with Accutase (Sigma-Aldrich) for 3 minutes at 37℃, quenched with growth media, and then counted. For each condition, 50,000 cells were used and blocked with Human TruStain FcX (Fc block, BioLegend) in FACS buffer (0.5% BSA (Sigma) in 1x PBS) for at least 15 minutes on ice, cells were kept on ice from this point forward. For flow cytometry in Figure 1, primary unconjugated antibodies including Mouse Isotype (Santa Cruz Biotechnology, sc-2025), Rabbit Isotype (Novus Biologicals, NB810-56910), anti-NCL (Santa Cruz Biotechnology, sc-17826), anti-hnRNP-U (Santa Cruz Biotechnology, sc-32315), anti-YBX1 (Santa Cruz Biotechnology, sc-101198), and anti-DDX21 (Novus Biologicals, NB100-1718) were bound in solution (precomplexed) to a Goat anti-Mouse AF647 secondary antibody (ThermoFisher Scientific, A32728) or a Goat anti-Rabbit AF647 secondary antibody (ThermoFisher Scientific, A27040) for at least 30 minutes on ice before using. The molar ratio was 2:1, primary:secondary. For flow cytometry in Figure 2, primary conjugated antibodies (AF647) were used directly for cell labeling and included mouse IgG1,k isotype (BioLegend, 400130), mouse IgG2b isotype (BioLegend, 400330) rabbit isotype (Novus Biologicals, NBP2-24891AF647), anti-NCL (Santa Cruz Biotechnology, sc-17826 AF647), anti-hnRNP-A1 (Santa Cruz Biotechnology, sc-32301 AF647), anti-hnRNP-U (Santa Cruz Biotechnology, sc-32315 AF647), anti-DDX21 (Novus Biologicals, NB100-1718AF647), anti-DNA-PKcs (Bioss Antibodies, BS-10994R), anti-PRDX1 (Santa Cruz Biotechnology, sc-137222 AF647), anti-TFRC (BioLegend, 334118 AF647), anti-STOM (Santa Cruz Biotechnology, sc-376869 AF647), anti-LAT1 (Santa Cruz Biotechnology, sc-374232 AF647), anti-GLUT1 (Santa Cruz Biotechnology, sc-58758 AF647), anti-EZR (Santa Cruz Biotechnology, sc-58758 AF647), anti-ꞵ2M (Santa Cruz Biotechnology, sc-13565 AF647), anti-GYPA (Santa Cruz Biotechnology, sc-53905 AF647). To the blocked cells, precomplexed antibody was added to a final concentration of 1 µg/mL (primary antibody) and allowed to bind to cells for 60 minutes on ice. After staining cells were spun at 4℃ for 3 minutes at 400x g and the supernatant discarded. All cell spins took place using these conditions. Cells were washed once with 150 µL of FACS buffer, spun under the same conditions, and finally resuspended in FACS buffer containing 0.1 µg/mL 4′,6-diamidino-2-phenylindole (DAPI, Sigma). Data collection occurred on a BD Biosciences LSRFortessa 3 and a gating strategy was used to isolate live, single cells, to examine antibody binding using FlowJo Software (FlowJo LLC).

To estimate antibody binding events per cell, we performed two additional measurements. Each of the AF647 primary conjugated antibodies listed above were analyzed for the protein (IgG) concentration and AF647 concentration to obtain a dye:IgG ratio. We also analyzed beads from the Quantum Alexa Fluor 647 MESF kit (Bang Labs) which have stocks of beads with specific numbers of AF647 dyes per bead. By building a standard curve of intensities from the flow cytometer using the beads we then calculated the number of dyes per cell and converted that to antibodies bound per cell. These measurements were all performed in biological triplicate. We reported the number of binding events per cell after subtracting the number of binding events we observed for the isotype antibody. In cases where a target antibody (e.g. NCL) had fewer binding events than the isotype for a given cell line (e.g. OCI-AML3), the reported number of binding events per cell was called zero.

### Confocal microscopy sample preparation

For adherent cells, culture conditions were similar to that described above however cells were deposited on glass coverslips #1.5 (Bioscience Tools, CSHP-No1.5-13) 24 hours before analysis. For suspension cells, culturing, counting, and Fc blocking were carried out similarly. Samples for “Live Cell” imaging (Figures 2, 3, 4, and 5) were processed as per the live cell flow cytometry noted above; however after staining and washing, cells were fixed with 3.7% formaldehyde (37% stock, Sigma) for 30 minutes at 25℃. Samples for “Fix/Perm” (Figure 2) were first fixed with 3.7% formaldehyde for 10 minutes at 25℃, rinsed once with 1x PBS, then permeabilized with 0.1% Triton X100 (Sigma) for 10 minutes at 25℃ and finally rinsed once with 1x PBS. Prior to staining, some experiments call for live cell RNase treatment. In these experiments, RNase A (Sigma) and ShortCut RNase III (New England Biolabs, NEB) would be added directly to the cell culture media for 2 hours before assay at a final concentration of 18 µM and 100 U/mL, respectively, supplemented with 2 mM MnCl_2_ (New England Biolabs, NEB). Control treated cells were incubated with a volume of 50% glycerol equivalent to the volume of RNase III added as well as 2 mM MnCl_2_. For staining reactions, primary unconjugated antibodies including Mouse Isotype (Santa Cruz Biotechnology, sc-2025), Rabbit Isotype (Novus Biologicals, NB810-56910), Goat Isotype (ThermoFisher Scientific, 02-6202), anti-hnRNP-U (ProteinTech, 14599-1-AP), anti-DDX21 (Novus Biologicals, NB100-1718), anti-ꞵ2M (Santa Cruz Biotechnology, sc-13565), anti-DNA-PKcs (Bethyl, A300-518A), and anti-dsRNA 9D5 (Absolute Antibody, Ab00458-23.0) were added to cells at 2.5 µg/mL final concentration for 45 minutes on ice. After staining, cells were washed twice with 1x PBS and then stained with a secondary antibody at a final concentration of 2.5 µg/mL targeting the primary antibody species with an appropriate fluorophore depending on the experiment: Goat anti-Mouse AF647 (ThermoFisher Scientific, A32728), Goat anti-Rabbit-AF647 (ThermoFisher Scientific, A27040), Donkey anti-Goat-AF647 (ThermoFisher Scientific, A-21447), Goat anti-Mouse AF568 (ThermoFisher Scientific, A-11031), and Goat anti-Rabbit-AF568 (ThermoFisher Scientific, A-11036). Secondary stains occurred for 30 minutes on ice and in the dark, after which cells were washed once with 1x PBS. A final fixation for the “Fix/Perm” samples was performed in parallel with the “Live Cell” samples with 3.7% formaldehyde in 1x PBS for 30 minutes at 25℃ in the dark. Nuclei were stained with 0.1 µg/mL DAPI in FACS buffer for 15 minutes at 25℃. Suspension cells were applied to glass slides using a CytoSpin centrifuge (Fisher Scientific): this was accomplished by centrifugation at 500x g for 5 minutes on a CytoSpin 1867. Finally, cells were mounted in ProLong Diamond Antifade Mountant (ThermoFisher Scientific) and a coverglass was sealed over the sample with nail polish. It should be noted that the incubation of live cells–especially suspension cells–on ice can result in substantial cell shrinkage relative to cells fixed prior to immunolabeling. All samples were then imaged on a Leica TCS SP8 STED ONE microscope.

### Confocal microscopy data acquisition and analysis

For all experiments, at least three regions of interest (ROIs) were acquired using a 63x oil immersion objective across one or more z-slices. Leica’s line-sequential scanning method was used and images were acquired at 1024 by 1024 resolution with a pinhole size of 1 AU. The DAPI channel was acquired with a PMT detector while all other channels were imaged using Hybrid detectors.

For colocalization analysis, ROIs comprising a sum of at least 20 cells were considered. These images were then processed using Imaris Microscopy Image Analysis software (Oxford Instruments). Here, a single z-slice from each region of interest was taken, selected to be near the middle of the cells (with respect to their z-thickness), and the spot-finder function was used to identify spots of roughly 0.5 µm. In-software background subtraction was used with default settings, and spots were selected by thresholding spot quality at the elbow of the distribution. This resulted in a series of x– and y-positions for each spot from each channel, which were then exported for quantitative analysis. Colocalization of spots from paired channels were analyzed by implementing a custom Python script (https://github.com/FlynnLab/jonperr) to identify the nearest neighbors of each spot (in nanometers, nm) with a k-d tree algorithm (scipy.spatial.KDTree). Then, the distances between nearest neighbors were calculated for each pair of targets across all ROIs and plotted in a histogram. To assess the relative fraction of each spot type (antibody #1) within the other pair’s spots (antibody #2), we calculated a Manders’ colocalization coefficient (MCC) using the aforementioned Python script. We performed this calculation in both directions: spots of Antibody #1 in total Antibody #2 spots, and the reverse.

To quantify and compare the intensities of spots on the cell surface across treatment conditions, Imaris was used to identify spots throughout the entire z-stack. Spots were again selected by thresholding for spots of quality greater than that associated with the elbow of the spot quality distribution. Then, the mean intensities of these distributions were saved as .xls files. Python script (https://github.com/FlynnLab/jonperr) was then used to quantitatively compare spot intensities across treatment groups. A nonparametric permutation test by bootstrapping was employed to determine the statistical significance of differences in spot intensity distributions. In this analysis, our null hypothesis stated that there was a greater than five percent chance that differences in medians between the union of two distributions would be greater than the observed difference. The test was performed with subsamples of size 100 over 10,000 iterations.

Intracellular fluorescence signal of TAT-eGFP, R5K-TAT-eGFP, and eGFP-treated cells was quantified first by using Imaris’ tool for cytoplasm and nucleus detection. The DAPI signal and eGFP signal were used to discriminate between the nuclei and cytoplasmic regions of cells. Nuclei of the A549 cells were estimated to be 12 um for this analysis. Inclusion of nuclei and cytoplasmic domains was thresholded by both thresholding at the elbow the “quality” distribution and excluding cells with nuclei with less than half of their total size in the frame. Next, the mean intensities of each cell were downloaded to .xls files. Intensities for each cell from replicates were concatenated and plotted in Prism. Prism was then used to calculate a Mann-Whitney U-test between cell intensity distributions and assess the statistical significance in differences in intensity between treatment conditions.

### Super resolution imaging and reconstruction

An overview of the method and processing can be found in Figure S3A. For single-molecule super-resolution microscopy we used direct stochastic optical reconstruction microscopy (dSTORM). To perform this, the PBS in which the cells were stored was replaced by a reducing oxygen scavenging buffer to induce blinking of fluorophores as described in literature (Halpern et al., 2015). The blinking buffer consisted of 2 μL/mL catalase (Sigma), 10% (w/v) glucose (BD Biosciences), 100 mM Tris-HCl (ThermoFisher Scientific), 560 μg/mL glucose oxidase (Sigma), and 20 mM cysteamine (Sigma). First, diffraction-limited images were obtained with low-intensity illumination of few W/cm^2^. Then, the laser power was increased to approximately 3 kW/cm^2^. Image acquisition was started after a short delay in order to ensure that most fluorophores were shelved into a dark state. The exposure time was 50 milliseconds (ms), and approximately 40000 frames were recorded.

As the data was obtained with a sCMOS camera, which typically exhibit few pixels with deviating sensitivity (“hot” and “cold” pixels), the obtained single-molecule data was corrected for individual pixels with abnormally high or low sensitivity first (Huang et al., 2011). 4000 frames of a raw data stack were averaged. Hot and cold pixels, which are a systematic deviation, persist, in contrast to the random single-molecule signals. Each pixel was compared to its neighbors, using 8-connectivity. If a deviation of more than 3% from the median of the neighboring pixel was observed, a correction factor that set the pixel to the median of its neighbors was recorded as previously described (Almahayni et al., 2023). Otherwise, the pixel was not considered to be significantly brighter or darker. This yielded a correction mask which was applied to all frames of the raw data. Finally, the no-light counts were subtracted from the pixel-corrected data.

For localization of single molecules, the Fiji plugin ThunderSTORM was used (Ovesný et al., 2014). Each camera frame was filtered with a B-spline filter of order 3 and scale 2. Local maxima, corresponding to single-molecule signals, were detected with 8-neighborhood connectivity and a threshold of 1.1 or 1.2 times the standard deviation of the first wavelet level. Detected local maxima were fitted with a 2D-Gaussian using least squares, and the position was recorded. Next, to account for single-molecule signals being active in multiple frames, merging of localizations was performed, using a maximum distance of 30 nm and maximum of 5 off frames with no limit regarding on frames. Cross correlation-based drift correction (magnification 5, bin size 5) was performed, followed by filtering of the localizations (sigma of the point spread function (PSF) between 60 and 270 nm, intensity below 37800 photons, localization uncertainty smaller than 30 nm). For visualization the final localizations were reconstructed as 2D histograms with magnification of 5 (corresponding to a pixel size of 17.7 nm).

For cluster analysis, an automated pipeline was established, using the raw list of localizations. Frist, Ripley’s H-function was calculated on three areas with a large number of well-separated clusters. The resulting preferential cluster size from the three areas (which was, notably, always very similar) was averaged, multiplied by a correction factor of 0.45, and used as the seed radius for the following DBscan analysis. This DBScan script yielded all individual clusters, the total number of clusters, the average number of points per cluster and the spatial relation between Clusters. The identified final clusters were then analyzed with respect to their spatial relation, size, and number of localizations. This unbiased approach follows recommended analysis procedures recently described (Nieves et al., 2023).

Crucially, each dataset was subjected to identical postprocessing and cluster analysis steps with no manual intervention, thus avoiding any biases arising from different parameter settings. Custom scripts used for this analysis can be shared upon request.

### Cell surface biotinylation, subcellular fractionation, and biotin capture and peptide generation

Cells (A549, HEK293, OCI-AML3, and K562) were cultured above and approximately 20 million cells were collected for processing; all in biological triplicate. Suspension cells were able to be processed directly in solution, while adherent cells were initially processed on the plates so as not to disturb the cell surface composition or plasma membrane integrity. Culture media was washed from cells twice with 1x PBS and then cell surface exposed proteins were labeled with sulfo-NHS-SS-biotin (APExBio) at a final concentration of 1 mg/mL in 1x PBS at 4℃ for 20 minutes. After labeling was complete, the reaction was quenched by adding 100mM Tris-HCl pH 8 (ThermoFisher Scientific) for 2 minutes at 4℃. This buffer was removed from cells and cells were washed once in fresh, ice cold 1x PBS. To obtain cytosolic and membrane fractions (Flynn et al., 2021), suspension cells were directly resuspended in Membrane Isolation Buffer (10mM HEPES (ThermoFisher Scientific), 250mM Sucrose (Sigma), 1mM EDTA) at 5 million cells per 1 mL. Adherent cells were collected off the plate by scraping in the ice cold PBS, pelleted, and then similarly resuspended at 5M cells / 1 mL of Membrane Isolation Buffer. Cells were rested on ice for up to 5 minutes, moved to a glass dounce homogenizer (Sigma), and then homogenized using 40-80 strokes to obtain a resuspension of approximately 50% released nuclei. Over douncing can cause nuclei rupture and contamination of the cytosolic fraction. After douncing, unbroken cells and nuclei were pelleted by spinning at 4℃ for 10 minutes at 700x g. Supernatants (cytosol and membranes) were carefully transferred to a new tube and pellets discarded. The supernatants were again centrifuged at 4℃ for 30 minutes at 10,000x g. Most (90%) of the supernatant was removed and saved as cytosolic fractions. The remaining supernatant was discarded as it was near to the membrane pellet. The membrane pellet was gently washed with 500 µL of ice cold 1x PBS (the pellet was not resuspended here), the tube spun down briefly, and all supernatant was discarded to leave a cleaned membrane pellet. Finally the membrane pellet was resuspended in 500 µL of CLIP lysis buffer (50 mM Tris-HCl pH 7.5, 200 mM NaCl (Sigma), 1 mM EDTA, 10% glycerol (ThermoFisher Scientific), 0.1% NP-40, 0.2% Triton-X100, 0.5% N-lauroylsarcosine). Both the cytosolic and membrane lysates were stored at –80℃ for later processing. This process was used both for western blotting (Figure 1) and Mass Spectrometry (Figure 2). In Figure 1, cytosolic and membrane pairs were loaded on an equivalent cell fraction basis to enable comparison on the relative distribution of proteins within a cell line, and normalized to the total mass of protein in the cytosolic fraction across cell lines.

For processing cytosolic and membrane fractions from A549, HEK293, OCI-AML3, and K562, lysates were first quantified with a BCA assay (ThermoFisher Scientific). A total of 30 µg of total protein from each sample was saved as input material. For biotin enrichment a total of 65 µg of the cytosol and 65 µg of membrane lysates were used, diluted into 350 µL of CLIP lysis buffer and captured with 33 µL of MagReSyn Streptavidin magnetic beads (Resyn Bio). The capture proceeded for 3 hours at 4℃ on rotation. After capture, beads were washed stringently using buffers headed to 37℃. The washes were all 1 mL per sample, and occurred in series: once with High Stringency buffer (15 mM Tris-HCl pH 7.5, 5 mM EDTA, 2.5 mM EGTA, 1% Triton-X100, 1% Sodium deoxycholate, 0.001% SDS (ThermoFisher Scientific), 1200 mM NaCl, 625 mM KCl), three times with 5% SDS in 0.5x PBS, and then once in 25℃ 1x PBS. Finally, leveraging the disulfide bond lability built into the sNHS-SS-biotin probe used the label the cell surface, biotinylated proteins were selectively eluted by incubating the beads in 100 µL of 5 mM dithiothreitol (DTT, Sigma), 5% SDS, in 1x PBS for 30 minutes at 75℃. After this cleavage step, the supernatant was moved to a new tube, the beads were rinsed once with pure water, and this water supernatant was pooled with the DTT-released proteins. Samples were frozen here for downstream processing.

To generate tryptic peptides, the S-trap column system was used (ProtiFi) as per the manufacturer’s protocol. Briefly, samples (input material and streptavidin enriched material) were thawed and SDS was added to a final concentration of 5% and 5 mM DTT was added fresh. Samples were denatured and reduced at 75℃ for 30 minutes after which iodoacetamide was added to a final concentration of 25 mM and incubated in the dark for 30 minutes at 25℃. Samples were then acidified by adding phosphoric acid to a final concentration of 1.2%, mixed by gently vortexing, and then the Bind/Wash Buffer (100 mM TEAB in 90% methanol) was added and again mixed by vortexing. Samples were then applied to the S-trap micro columns with centrifugation (all spins were for 10 seconds and 400x g). Samples were washed three times with Bind/Wash Buffer. To the washed, packed column (now loaded with the protein sample) 20 µL of Trypsin digestion solution (1 µg MS-grade Trypsin (Promega) in 50 mM ammonium bicarbonate) was added and samples were incubated at 47℃ for 2 hours. Digested tryptic peptides were spun out of the column and residual peptides were eluted by sequentially adding 40 µL of 0.2% formic acid in water and then 50% acetonitrile in water. The three elutions were pooled and dried using a SpeedVac and then lyophilizer before analyzing via mass spectrometry. The peptides were resuspended in 20% acetonitrile in water containing 0.1% formic acid and bath sonicated for 5 minutes. These resuspended peptides were centrifuged for 10 minutes at 20,000x g and then transferred to mass spectrometry vials for injection and analysis.

### Mass spectrometry data generation

For all peptides generated, we followed the same procedure for mass spectrometry analysis and peptide database searching. Specifically, a nanoElute was attached in line to a timsTOF Pro equipped with a CaptiveSpray Source (Bruker, Hamburg, Germany). Chromatography was conducted at 40℃ through a 25 cm reversed-phase C18 column (PepSep) at a constant flow rate of 0.5 µL/minute. Mobile phase A was 98/2/0.1% water/acetonitrile/formic acid (v/v/v) and phase B was acetonitrile with 0.1% formic acid (v/v). During a 108 minute method, peptides were separated by a 3-step linear gradient (5% to 30% B over 90 minutes, 30% to 35% B over 10 minutes, 35% to 95% B over 4 minutes) followed by a 4 minute isocratic flush at 95% for 4 minutes before washing and a return to low organic conditions. Experiments were run as data-dependent acquisitions with ion mobility activated in PASEF mode. MS and MS/MS spectra were collected with *m*/*z* 100 to 1700 and ions with *z* = +1 were excluded. Raw data files were searched using PEAKS Online Xpro 1.7 (Bioinformatics Solutions Inc., Waterloo, Ontario, Canada). The precursor mass error tolerance and fragment mass error tolerance were set to 20 ppm and 0.03, respectively. The trypsin digest mode was set to semi-specific and missed cleavages was set to 3. The human Swiss-Prot reviewed (canonical) database version 2020_05 (downloaded from UniProt) and the common repository of adventitious proteins (cRAP v1.0, downloaded from The Global Proteome Machine Organization) totaling 20,487 entries were used. Carbamidomethylation was selected as a fixed modification. Oxidation (M), Deamidation (NQ) and Acetylation (N-term) were selected as variable modifications. Raw data files and searched datasets are available on the Mass Spectrometry Interactive Virtual Environment (MassIVE), a full member of the ProteomeXchange consortium under the identifier: MSV000092005. The complete searched datasets are also available in our supplementary files.

### Mass spectrometry data analysis

For the analysis of MS database results from cell-surface proteomics performed with sulfo-NHS-SS-biotin, any proteins detected from species other than human were purged from the dataset in Excel. Then, all keratins were removed from the resulting list of proteins to minimize the presence of contaminants from desquamation during peptide processing. Hits from replicates of each fraction were concatenated in Python, and the peak areas associated with hits in each fraction were plotted against their abundance rank as determined by peak area. Scipy’s curve_fit was used to define an exponential line of best fit in the form y = b+ a/(x^c). The derivative of the line of best fit was taken, and the hits were thresholded at the y-value (peak area) where this derivative is equal to –1. Only proteins with peak areas above the designated threshold for all three replicates were retained in the list of candidate hits. Next, proteins with fewer than 3 unique peptides were filtered out using the same Python script. The list of proteins obtained through this process were defined as hits.

To perform GO-term analysis, all proteins observed across all four fractions from each cell line were concatenated, creating a list of background proteins from each of the four cell lines tested. Lists of hits from the membrane hits pulldown from each cell line were submitted to DAVID (https://david.ncifcrf.gov/) and ran against their respective backgrounds. Then, for each category of GO-term (BP, CC, and MF), the union of the top four terms across all cell lines and their associated sizes and Benjamini values was plotted for comparison of enrichment across cell lines.

### Cell surface proximity labeling of proteins, peptide generation, and data analysis

Samples were prepared similarly to the flow cytometry workflow as described above however rather than dye-conjugated secondaries, here horseradish peroxidase (HRP) conjugates secondaries were used. The isotype (control) or anti-RBP (target) primary unconjugated antibody (listed above) was precomplexed with an appropriate secondary-HRP antibody at 2:1, primary:secondary for at least 30 minutes on ice. Cells were grown as biological triplicate cultures and typically 2.5 million cells were used per replicate per labeling experiment. Cells were harvested from culture, washed of culture media, and resuspended in ice cold FACS buffer to which the Fc blocker was added for at least 15 minutes. After blocking cells were adjusted to 1 million cells per mL of FACS buffer and then the precomplexed antibodies were added for staining to a final concentration of 2.5 µg/mL. Staining occurred for 60 minutes at 4℃ on rotation, after which cells were pelleted, supernatants discarded and cells washed once in ice cold 1x PBS. This wash is important to remove excess BSA in the FACS buffer. Next, cells were gently but quickly resuspended in 980 µL of 100 µM biotin-phenol (Iris Biotech) in 1x PBS at 25℃. To this, 10 µL of 100 mM H_2_O_2_ was quickly added, tubes capped and inverted, and the reaction allowed to proceed for 2 minutes at 25℃. Precisely after 2 minutes, the samples were quenched by adding sodium azide and sodium ascorbate to a final concentration of 5 mM and 10 mM, respectively. Samples were inverted and cells pelleted at 4℃. Samples were then washed sequentially once with ice cold FACS buffer and then twice ice cold 1x PBS, after which cell pellets were lysed in 500 µL of CLIP lysis buffer, briefly sonicated to solubilize chromatin, and frozen at –80℃ for later processing.

Once all the proximity labeling was complete, lysates were thawed in batches to perform the following steps in parallel. Total protein amounts were quantified using the BCA assay and labeling efficiency and consistent was checked using Western blotting. For biotin enrichment, we used the streptavidin western QC to determine the biotin signal across all samples first, then calculated the total µg of lysate that was needed from that sample to generate 5,000,000 units of streptavidin-IR800 signal on the LiCor. That µg value was then used as input mass for each of the replicates across all of the proximity labeling samples for the biotin capture and MS prep. Samples were normalized to a final volume of 500 µL with CLIP lysis buffer and to each sample 100 µL of Pierce NeutrAvidin Agarose (ThermoFisher Scientific) was added and incubated at 4℃ for 4 hours on rotation. Beads were then washed twice with 1 mL of High Stringency, twice with 1 mL of 4 M NaCl in 100 mM HEPES all at 37℃. Salts and detergents were then rinsed from the beads by sequentially washing twice with 1 mL 1x PBS, twice with 1 mL of LC-MS grade water (Fisher Scientific) and finally with 1 mL of 50 mM ammonium bicarbonate. An on-bead trypsin digestion was then set up by adding 200 µL of 50 mM ammonium bicarbonate and 1 µg MS-grade Trypsin to each sample and incubating for 16 hours at 37℃ with occasional shaking. After digestion, the samples were acidified by adding formic acid to a final concentration of 0.5%. The solution containing released peptides was moved to a new tube and the beads were rinsed twice with 300 µL LC-MS grade water to capture any remaining peptides. All elution and wash samples for a given replicate were combined and SpeedVac’ed to a volume of < 200 µL. Samples were then desalted using a C18 spin column (ThermoFisher Scientific): preconditioned with 50% methanol in water, washed twice with 5% acetonitrile, 0.5% formic acid in water, the sample bound twice to the column, then the columns washed four times with 0.5% formic acid in water. Finally the peptides were eluted into Protein LoBind tubes (Eppendorf) with two applications of 40 µL of 70% acetonitrile, 0.5% formic acid in water. Organic solvents were removed with a SpeedVac and samples were fully dried with a lyophilizer. Resulting peptides were analyzed on the timsTOF Pro as described above.

Data generated from proximity labeling samples were searched against the reference proteomes as noted above. To identify enriched proteins from these datasets we took an approach that broadly compared enrichment in the target antibody experiments (e.g. csRBPs) to that of control antibodies (e.g. Isotypes). Results of the database search were first purged of non-human proteins and keratins as previously described. All proteins with fewer than two unique peptides were also filtered out of the list. Next, Excel was used to calculate the mean spectral count for each of the remaining proteins across triplicates. For each protein, the mean spectral counts associated with each antibody probe were divided by the mean spectral counts of the corresponding isotype, creating an enrichment factor of each protein in the proximity labeling over the isotype. Python was then used to calculate the ratio of total protein isolated from proximity labeling with protein-targeting antibodies divided by that collected with isotype antibodies. Proteins from each set of proximity labeling data with enrichments either less than two or the previously described ratio were filtered from the dataset. The resultant lists comprise the hits associated with each round of proximity labeling.

### Cell surface proximity labeling of RNAs and gel analysis

Samples were prepared here in a similar manner as described above for the proximity label of proteins with key differences. The HRP-conjugated secondary antibody was switched for Protein A-HRP (Cell Signaling) and used at the same molar ration; 2 primary antibodies per 1 Protein A-HRP molecule. The biotinylation reagent used here was biotin-aniline (Iris Biotech), it was used at 200 µM final concentration in 1x PBS for the labeling reaction, and the labeling reaction was allowed to proceed for 3 minutes at 25℃. After pelleting the cells from the quenching reaction, cells were directly lysed and total RNA isolated as described before (Hemberger et al., 2023). Briefly, RNAzol RT (Molecular Research Center, Inc.) was used to lyse cell pellets by placing the samples at 50℃ and shaking for 5 min. To phase separate the RNA, 0.4X volumes of water was added, vortexed, let to stand for 5 minutes at 25℃ and lastly spun at 12,000x g at 4℃ for 15 min. The aqueous phase was transferred to clean tubes and 1.1X volumes of isopropanol was added. The RNA is then purified over a Zymo column (Zymo Research). For all column cleanups, we followed the following protocol. First, 350 μL of pure water was added to each column and spun at 10,000x g for 30 seconds, and the flowthrough was discarded. Next, precipitated RNA from the RNAzol RT extraction (or binding buffer precipitated RNA, below) is added to the columns, spun at 10,000x g for 10-20 seconds, and the flowthrough is discarded. This step is repeated until all the precipitated RNA is passed over the column once. Next, the column is washed three times total: once using 400 μL RNA Prep Buffer (3M GuHCl in 80% EtOH), twice with 400 μL 80% ethanol. The first two spins are at 10,000x g for 20 seconds, the last for 30 seconds. The RNA is then treated with Proteinase K (Ambion) on the column. Proteinase K is diluted 1:19 in water and added directly to the column matrix and then allowed to incubate on the column at 37℃ for 45 min. The column top is sealed with either a cap or parafilm to avoid evaporation. After the digestion, the columns are brought to room temperature for 5 min; lowering the temperature is important before proceeding. Next, eluted RNA is spun out into fresh tubes and a second elution with water is performed. To the eluate, 1.5 μg of the mucinase StcE (Sigma-Aldrich) is added for every 50 μL of RNA, and placed at 37℃ for 30 minutes to digest. The RNA is then cleaned up again using a Zymo column. Here, 2X RNA Binding Buffer (Zymo Research) was added and vortexed for 10 seconds, and then 2X (samples + buffer) of 100% ethanol was added and vortexed for 10 seconds. The final RNA is quantified using a Nanodrop. In vitro RNase or Sialidase digestions took place by digesting 50 μg total RNA with either, nothing, 4 μL RNase Cocktail (ThermoFisher Scientific), or 4 μL of α2-3,6,8,9 Neuraminidase A (NEB) in 1x NEB Glyco Buffer #1 (NEB) for 60 minutes at 37℃. After digestion, RNA was purified using a Zymo column as noted above and was then ready for gel analysis.

In order to visualize the labeled RNA, it is run on a denaturing agarose gel, transferred to a nitrocellulose (NC) membrane, and stained with streptavidin (Hemberger et al., 2023). After elution from the column as described above, the RNA is combined with 12 μL of Gel Loading Buffer II (GLBII, 95% formamide (ThermoFisher Scientific), 18 mM EDTA (ThermoFisher Scientific), 0.025% SDS) with a final concentration of 1X SybrGold (ThermoFisher Scientific) and denatured at 55℃ for 10 minutes. It is important to not use GLBII with dyes. Immediately after this incubation, the RNA is placed on ice for at least 2 minutes. The samples are then loaded into a 1% agarose, 0.75% formaldehyde, 1.5x MOPS Buffer (Lonza) denaturing gel. Precise and consistent pouring of these gels is critical to ensure a similar thickness of the gel for accurate transfer conditions; we aim for approximately 1 cm thick of solidified gel. RNA is electrophoresed in 1x MOPS at 115V for between 34 or 45 min, depending on the length of the gel. Subsequently, the RNA is visualized on a UV gel imager, and excess gel is cut away; leaving ∼0.75 cm of gel around the outer edges of sample lanes will improve transfer accuracy. The RNA is transferred with 3M NaCl pH 1 (with HCl) to an NC membrane for 90 minutes at 25℃. Post transfer, the membrane is rinsed in 1x PBS and dried on Whatman Paper (GE Healthcare). Dried membranes are rehydrated in Intercept Protein-Free Blocking Buffer, TBS (Li-Cor Biosciences), for 30 minutes at 25℃. After the blocking, the membranes are stained using streptavidin-IR800 (Li-Cor Biosciences) diluted 1:5,000 in Intercept blocking buffer for 30 minutes at 25℃. Excess streptavidin-IR800 was washed from the membranes using three washes with 0.1% Tween-20 (Sigma) in 1x PBS for 3 minutes each at 25℃. The membranes were then briefly rinsed with PBS to remove the Tween-20 before scanning. Membranes were scanned on a Li-Cor Odyssey CLx scanner (Li-Cor Biosciences).

### Immunoprecipitation and western blotting

HeLa cells stably expressing DDX21-HA (Calo et al., 2015) were cultured as above. Cell surface protein labeling was accomplished using sulfo-NHS-biotin (APExBio, no disulfide bond) as described above after which crude membrane fractions were isolated as described for the adherent cells. After isolation and solubilization with CLIP lysis buffer, total protein quantification occurred using the BCA assay. For each sample 5 μg of protein was used for input and 20 μg used for either anti-HA coated bead (ThermoFisher Scientific) or streptavidin coated bead (MyOne C1 beads, ThermoFisher Scientific) enrichments. In both cases, 10 μL bead surrey was added to the membrane lysates in 100 μL of CLIP lysis buffer and binding occurred at 4℃ for 16 hours. After binding, the beads were washed three times with High Stringency buffer and then twice with 1x PBS. Proteins were released from the beads by heating at 85℃ for 10 minutes in 20 µL 5 mM DTT and 1x LDS (ThermoFisher Scientific) and 1 mM free biotin (ThermoFisher Scientific). Input and enriched proteins were analyzed by Western blot, staining with anti-HA (Santa Cruz Biotechnology, sc-7392) and streptavidin-IR800 and finally scanning on Li-Cor Odyssey CLx scanner.

### Recombinant protein expression and cell treatments

To produce eGFP-TEV-6xHIS, TAT-WT-eGFP-TEV-6xHIS, and TAT-R5K-eGFP-TEV-6xHIS proteins we used the Juice Bulk E. coli Cell-Free expression kit from Liberum Biotech Inc. as per the manufacturer’s protocol. In brief, plasmids for each three proteins were designed (**Table S5**). Plasmids were subcultured with NEB 5-alpha Competent E. coli, High Efficiency, (NEB) with Ampicillin (Sigma) and DNA isolated using a Plasmid MiniPrep Kit (Zymo Research). Protein expression runs were performed in 1 mL reaction volumes with a final concentration of 15 nM plasmid DNA for each reaction, which were left overnight at 26℃ in the dark in a Feather cartridge (Liberum Biotech Inc.). After 16 hours, the lysate was spun out of the Feather cartridge at 4000x g and added 100 µL of Ni-NTA agarose resin (Qiagen) to capture HIS-tagged proteins. Lysate was incubated at 25℃ for 30 minutes on rotation, after which the resin was washed three times with 250 µL Wash Buffer (250 mM NaCl (Sigma), 20 mM imidazole (Sigma), 5% glycerol (Sigma), 50 mM HEPES (Gibco), pH 7.5, 0.5 mM TCEP (Sigma)). HIS-tagged proteins were eluted in 2×100 µL of Elution Buffer (250 mM NaCl, 250 mM imidazole, 5% glycerol, 50 mM HEPES, pH 7.5, 0.5 mM TCEP). Total protein mass was estimated using the Pierce BCA Protein Assay Kit (ThermoFisher Scientific) and subsequently the 6xHIS tag was cleaved using the TEV Protease (NEB) by adding TEV enzyme assuming all protein isolate was 6xHIS tagged based on the activity of the TEV protease. TEV cleavage was allowed to proceed under the manufacturer’s protocol for 60 minutes at 30℃. Excess 6xHIS tag and TEV protease enzyme were removed by adding 100 µL of Ni-NTA resin and captured as per above. The unbound portion of this capture step was kept as HIS-free eGFP protein and conjugates and concentrated once on a 10K Amicon column (Millipore Sigma) by spinning for 6-8 minutes at 6000x g. The protein was washed once by adding 400 µL of 1x PBS to the top of the Amicon column and spinning again, the finally concentrated protein quantified as above. We observed that TAT-WT and TAT-R5K eGFP conjugates (but not eGFP alone) were contaminated with nucleic acids (A260nm signal on the Nanodrop (ThermoFisher Scientific) from the expression while eGFP alone was not. To address this we performed a nuclease treatment of all three proteins after TEV cleavage to destroy any carry-over E. coli RNA or DNA by adding 1 µL RNase Cocktail (ThermoFisher Scientific) and 1U TurboDNase (ThermoFisher Scientific) per 200 µg purified protein. These nuclease-digested samples were then cleaned using a new 10K Amicon column and quantified as described above. Purity and TEV cleavage were assessed using total protein blotting.

To characterize the internalization of eGFP fusion proteins, 10,000 A549 cells were plated in each well of a 8-chambered coverglass system (Cellvis, C8-1.5H-N) and cultured as above for 24 hours. Live cell RNase treatment was performed as noted above for the appropriate samplesFollowing the 2 hour incubation, all cells were washed with conditioned media (RNase-free) before adding conditioned media with 20 µg/mL eGFP fusion proteins and allowed to incubate at 37℃ for 20 minutes in the growth incubator. After incubating, cells were washed twice with 1x PBS before fixation with 3.7% formaldehyde in 1x PBS for 30 minutes at 25℃ in the dark. Nuclei were stained with DAPI as mentioned above, and the GFP channel was imaged by confocal microscopy as described above. For assessment of cell surface binding, after no treatment or RNase digestion and 1x PBS washing, cells were cooled on ice and 20 µg/mL eGFP fusion proteins were added in FACS buffer. After binding, cells were washed twice with 1x FACS buffer before Fc blocking and anti-GFP staining was performed as per the cell surface staining protocol above. A rabbit polyclonal anti-GFP antibody (ThermoFischer, A11122) and AlexaFluor 568-conjugated goat anti-rabbit antibody (ThermoFischer, A11036) was used to visualize cell surface eGFP fusion proteins.

### Analysis of publicly available datasets

#### RNA binding proteins

RBPomes were compiled by extracting 48 lists of RBPomes from RBPbase (https://rbpbase.shiny.embl.de/). 30 of these RBPomes were sourced from experiments and 18 were isolated from lists of gene annotations. An Excel was used to intersect the GENCODE names for genes corresponding to RBPs. The RBP ID was plotted in a histogram displaying the number of datasets that RBP was identified in, generating values between one and 36 hits. The threshold of 11 (RBP found in 11 or more datasets) was chosen, as this point represents the elbow of the plot, minimizing the inclusion of proteins rarely observed to bind RNA. Thresholding such that a protein defined as an RBP must be observed in at least 11 RBPomes yielded a list of 1072 RNA-binding proteins.

#### Surfaceomes

Surfaceomes were collected from 23 publicly available peer-reviewed works and converted into their GENCODE names. Datasets were classified by the chemistry used to selectively label cell-surface proteins: sulfo-NHS-SS-biotin, periodate, or a collection of miscellaneous methods. Within each of these chemistries, Python script was used to count the number of times each RBP appeared. All RBPs appearing at least once in a chemistry were added to a list corresponding to sulfo-NHS-SS-biotin, periodate, or the group of miscellaneous methods. Each of these lists was intersected to generate a list of RBPs identified by all three chemistries. Then, using a histogram of the number of datasets each of these RBPs appeared in, a threshold was set near the shoulder of the left tail of the distribution, requiring that an RBP to be observed in at least five datasets. This list was defined as the csRBP candidate list.

## Supporting information

Table S1. RBPome and cell-surface proteome analysis.

Table S2. Cell-surface proteomics with sulfo-NHS-SS-biotin and subcellular fractionation.

Table S3. dSTORM image settings and quantitation.

Table S4. Cell-surface proximity labeling proteomics.

Table S5. Plasmid sequences for cell-free expression of eGFP and TAT-eGFP fusion proteins.

## Acknowledgments

We thank Phillip A. Sharp, Kayvon Pedram, Julia Belk, Brian Do, Robert C. Spitale, Carolyn R. Bertozzi and other members of the Flynn Lab for helpful comments and discussions. This work was supported by grants from Burroughs Wellcome Fund Career Award for Medical Scientists (R.A.F.), the Sontag Foundation Distinguished Scientist Award (R.A.F.), the Rita Allen Foundation (R.A.F.), a private donation administered by the National Philanthropic Trust (R.A.F.), the Beckman Young Investigator from the Arnold and Mabel Beckman Foundation (B.Z.), a grant from the Else-Kröner-Fresenius-Stiftung (2020_EKEA.91) and by the Max Planck Society (K.A., G.N., M.S., and L.M.).

## Author Contributions

R.A.F. conceived and supervised the project. N.B., J.P., and R.A.F. performed data analysis on publicly available RBP and surface proteomics datasets. R.A.F. and J.P. performed most of the cell culture, and confocal imaging, TAT addition assays, and data analysis, with help from P.C., C.G.L., R.M.C., and H.H. Mass Spec data collection and analysis strategies were developed by A.L., R.V., and B.Z. Cell culture, light microscopy, and super resolution reconstruction and data analysis was performed by K.A., G.N., M.S., and L.M., E.C., and K.T. provided cells and performed experimental design. R.A.F. and J.P. wrote the manuscript. All authors discussed the results and revised the manuscript.

